# Evidence of multiple origins of glyphosate resistance evolution in *Lolium multiflorum*

**DOI:** 10.1101/2021.06.24.449792

**Authors:** Caio A. C. G. Brunharo, Matthew A. Streisfeld

## Abstract

- The multitude of herbicide resistance patterns in the weed *Lolium multiflorum* L. is a remarkable example of the rapid adaptation to anthropogenic-driven disturbance. Recently, resistance to glyphosate was identified in multiple populations of *L. multiflorum* in Oregon.
- We used phenotypic approaches, as well as population genomic and gene expression analyses, to determine if known mechanisms were responsible for glyphosate resistance, if resistance phenotypes evolved independently in different populations, and to identify potential loci contributing to resistance.
- We found no evidence of genetic alterations or expression changes at known target and non-target sites of glyphosate resistance. Population genomic analyses indicated that resistant populations tended to have largely distinct ancestry from one another, with little evidence of admixture, suggesting that glyphosate resistance did not spread among populations via gene flow. Rather, resistance appears to have evolved independently on different genetic backgrounds. We also detected potential loci associated with the resistance phenotype, some of which encode proteins with potential effects on herbicide metabolism.
- Our results suggest that Oregon populations of *L. multiflorum* evolved resistance to glyphosate due to a novel mechanism. Future studies that characterize the gene or genes involved in resistance will be necessary to confirm this conclusion.

## INTRODUCTION

The human population is expected to reach nearly 10 billion by 2050, and meeting agricultural demands remains one of the biggest challenges for our society (United Nations, 2019). Weed interference can significantly reduce crop yields (Oerke, 2006). For example, wheat exhibits a 39-62% reduction in yield when infested with the agricultural weed *Lolium multiflorum* L. (Appleby et al., 1976). Weeds of agricultural crops are primarily managed with herbicides, as other techniques are less efficient and typically more expensive. The overreliance on herbicides as the main weed management tool in agriculture has selected for herbicide resistant weed populations. To date, 514 examples of herbicide resistance have been reported (Heap, 2021) from over 90 different crops and 70 countries around the world. Herbicide resistance poses a serious challenge for sustainable weed management, because new herbicides have not been developed for marketing in recent years, and there are numerous additional costs associated with non-chemical control methods. Furthermore, in some situations, fields with a long history of no-tilling that contain herbicide resistant weeds may have to return to conventional weed control techniques, increasing the carbon footprint of food production and moving against basic concepts of sustainable agriculture (Pretty, 2018).

The rapid and repeated evolution of herbicide resistance in agricultural fields is a clear example of parallel evolution due to the presence of similar selective pressures (Bolnick et al., 2018). There are many examples where the same genetic change resulted in the evolution of herbicide resistance across distinct plant lineages. To illustrate, over 90 populations from 40 different weed species have a mutation at position 197 of acetolactate synthase (ALS), conferring resistance to different herbicides that inhibit this enzyme (Heap, 2021). Despite this clear convergence on a single mutation site, the genetic basis of herbicide resistance may involve multiple mechanisms: mutations can occur in the target-site of the herbicide (denominated target-site resistance, TSR) or elsewhere (non-target-site resistance, NTSR) (Baucom, 2019). TSR has been demonstrated in several plant systems, and the functional basis of these mechanisms has been widely elucidated. A well-studied example comes from mutations in the gene that encodes enolpyruvylshikimate-3-phosphate synthase (EPSPS). This is a key enzyme in the shikimate pathway that is crucial for aromatic amino acid synthesis. The activity of EPSPS is inhibited by the herbicide glyphosate at the phosphoenolpyruvate binding site (Steinrücken & Amrhein, 1984). Amino acid substitutions in the active site of the enzyme, specifically at position 106, have been shown to provide reduced glyphosate binding due to structural conformation changes (Funke et al., 2006). This mutation does not dramatically reduce the catalytic efficiency of EPSPS in the presence of glyphosate, allowing treated plants to survive. TSR may also be conferred by deletions in the herbicide target-site, as is commonly found for protoporphyrinogen oxidase inhibitors (Patzoldt et al., 2006), or because of duplication of the target-site gene (Gaines et al., 2010). In the latter scenario, increased target-site expression is conferred by a higher concentration of the enzyme in the plant cells, which in turn requires a substantially higher concentration of herbicide to be absorbed, which is often impractical.

NTSR mechanisms are classified as those not involved in the target-site. The physiological bases of these mechanisms have been broadly described. In general, weeds with NTSR may exhibit reduced herbicide uptake, reduced translocation to the site of action, and enhanced herbicide degradation (reviewed by Delye et al., 2013). In recent years, many researchers have focused on elucidating the pathways involved in herbicide degradation (i.e. metabolic resistance), and results converge toward the involvement of cytochrome P450s, glutathione S-transferases, and membrane transporters (e.g. ATP-binding cassette transporter; ABC transporter). Most conclusions, however, come from indirect evidence, such as the application of P450 inhibitors followed by herbicide treatment (Oliveira et al., 2018), which would reverse resistance if enhanced activity of P450 is responsible for the resistance phenotype. However, the genes involved in resistance are still largely unknown (Han et al., 2021). The primary reason for the recent increased interest in metabolic herbicide resistance is the observation that weed populations with these types of resistance mechanisms commonly exhibit resistance to multiple herbicides from different chemical groups and mechanisms of action (Dimaano et al., 2020). Therefore, NTSR may confer resistance to weed populations against herbicides they have never been exposed to (Busi & Powles, 2016).

Although TSR is believed to be conferred by single major-effect alleles and likely contributes to much of the parallelism in resistance phenotypes among weed species, NTSR is believed to be a quantitative trait, conferred by multiple loci of small effect (Delye, 2013; Delye et al., 2013). However, recent research has shown that the resistance phenotype can be explained by major-effect genes, such as enhanced expression of the P450 *CYP81A10v7* that conferred resistance to seven herbicide chemistries (Han et al., 2021). While physiological modifications that lead to NTSR in weeds are well documented, the genetic basis of the modifications are poorly understood (Suzukawa et al., 2021).

*Lolium multiflorum* is a diploid (*2n* = *2x* = 14), obligate outcrossing winter annual ryegrass species of broad occurrence in the United States (DiTomaso & Healy, 2007) and throughout the world. It is native to the Mediterranean basin, and because of its desirable forage characteristics, this species has been adopted as a crop in many regions of the world (Humphreys et al., 2010). Although *L. multiflorum* is cultivated as a crop, this species may also be considered a weed when it grows where it is not desired. For instance, commercial *L. multiflorum* varieties are often sown as cover crops in the USA to enhance soil health indices in corn-soybean rotations (Shipley et al., 1992). However, persistence of *L. multiflorum* in subsequent growing seasons is not desirable, because it could compete with the cash crop early in the season. For clarity, hereafter, we refer to “annual ryegrass” when discussing the crop, and *L. multiflorum*, when describing the weed. The cultivated varieties of annual ryegrass, although the same species as the weedy biotypes, exhibit desirable traits associated with yield, including high nitrogen content, high seed vigor, and they are susceptible to herbicides. The weedy populations display undesirable characteristics as a result of the uncontrolled crossing outside of breeding programs and a continued process of de-domestication.

*L. multiflorum* has evolved resistance to many herbicides (Suzukawa et al., 2021). For example, a population collected from a prune orchard in California exhibited resistance to four different mechanisms of action (Brunharo & Hanson, 2018). In Oregon, *L. multiflorum* is a weed in many crops, including perennial ryegrass, tall fescue, wheat, orchardgrass, and annual ryegrass grown for seed. Recently, 60 resistant *L. multiflorum* populations were identified in Oregon, with some populations resistant to up to four different herbicide mechanisms of action (Bobadilla et al., 2021). The widespread herbicide resistance in grass seed fields in Oregon poses a serious threat to local agricultural communities, because of potential contamination of grass seed lots with herbicide resistant weed seeds. The seed lots are sold for many uses nationally and internationally, posing a potential source for dispersal of herbicide resistance genes.

Basic knowledge of weed adaptation to herbicides, dispersal, and detection are of primary importance to initiate a mitigation plan and to contain expansion of the areas infested with herbicide resistance. Gene flow from herbicide resistant weed populations into annual ryegrass crops is deemed as one of the main concerns farmers have, and its spread is commonly attributed to movement of agricultural machinery, commodity movement, and livestock feed (Schroeder et al., 2018). Understanding the genetic relatedness among herbicide resistant and susceptible weed populations may give clues to potential mechanisms of propagule dispersal. For example, if spread of a resistant weed population is through seed lot contamination, then policy makers could implement new rules or recommend new practices to prevent seed lot contamination (i.e. recommend seed certification based on herbicide resistance testing of seed lots, recommend longer rotations in the field, and others).

In herbicide resistant populations of *L. multiflorum* in Oregon, little is known about the underlying genetic mechanisms conferring herbicide resistance, or the genetic relationship among populations. In this study, we analyze patterns of genetic variation and admixture among 16 of these Oregon populations that vary in their resistance to glyphosate. The three primary objectives of this work were to 1) determine if resistance is conferred by known resistance mechanisms, 2) determine if resistance phenotypes evolved independently in different populations, and 3) identify potential loci involved in glyphosate resistance.

## MATERIALS AND METHODS

### Study populations

A set of 16 *L. multiflorum* populations from agricultural fields in the Willamette Valley in Oregon were identified for this study (Figure 1, Supporting Information Table S1). The populations were collected in 2017-2018 as part of a broader survey of herbicide resistance (Bobadilla et al., 2021). A cultivated, public variety of annual ryegrass known as “Gulf’ and a previously characterized multiple-herbicide resistant *L. multiflorum* population called PRHC from California (Brunharo & Hanson, 2018) were included in the study, as was a cultivated variety of perennial ryegrass (*L. perenne*) used as an outgroup. Gulf has been widely used as a reference susceptible population for *L. multiflorum* herbicide resistance characterization (Bobadilla et al., 2021), whereas PRHC is a population that exhibits resistance to four different herbicide mechanisms of action, including a known target-site mutation in *EPSPS*. The populations selected exhibited various herbicide resistance patterns, where resistance to ALS (mesosulfuron and pyroxsulam), acetyl-CoA-carboxylase (ACCase; clethodim, pinoxaden, and quizalofop), and EPSPS (glyphosate) inhibitors were the most common. Populations susceptible to all herbicides tested were also included in the study. However, in this study, we focused only on glyphosate because of its importance in agricultural and non-cropping areas, and because the NTSR mechanisms of resistance are largely unknown.

**Fig. 1.**
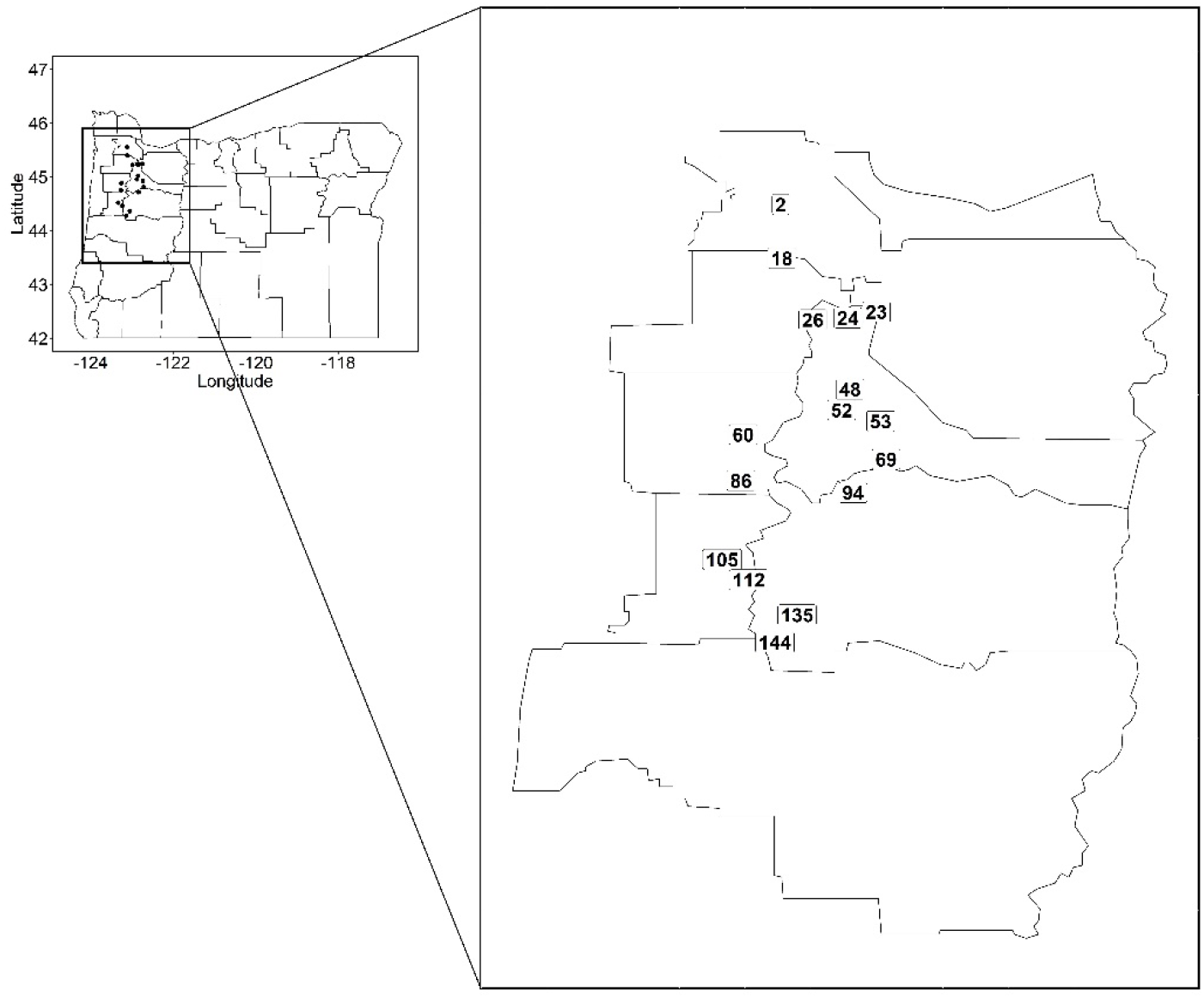
Collection sites of the 16 *Lolium multiflorum* populations used in this study.

### Phenotyping

To confirm glyphosate resistance in the sampled populations, we performed a shikimate accumulation assay. This method was implemented as a biomarker for glyphosate effects, because glyphosate inhibits EPSPS in the shikimate pathway, resulting in the accumulation of shikimate in the tissues (Dayan et al., 2015). Because this assay is quantitative, we were able to accurately diagnose the level of glyphosate resistance in each sampled individual across populations. If shikimate accumulation levels are distributed continuously within and among populations, then it would suggest quantitative control of glyphosate resistance, likely due to the contribution of multiple genetic loci. Conversely, qualitative levels of shikimate accumulation likely indicate a simple genetic basis for glyphosate resistance. Field collected seeds were germinated in petri dishes, and seedlings were transplanted to 0.5 L pots filled with commercial potting mix and grown in a greenhouse at 24 C and 14/10h (day/night). Twenty plants were grown from each *L. multiflorum* population. When plants reached the 23-BBCH (three tillers) growth stage (Hess et al., 1997), leaf tissue from the second youngest fully expanded leaves was collected, frozen in liquid nitrogen, and stored in a −80 C freezer until further analysis. After tissue sampling, glyphosate was applied at 1456 g acid equivalent per hectare (g a.e. ha^-1^).

Forty-eight hours after glyphosate application, the youngest fully expanded leaves from a different tiller were collected, weighed, and stored in Eppendorf tubes in a −80 C freezer until shikimate accumulation quantification was performed as described in Shaner et al. (2005). Briefly, samples were pulverized in liquid nitrogen, and 1000 μL of 10 mM ammonium phosphate monobasic (0.1% Tween, pH 4.4 with 0.1 HCl or NaOH) were added to the ground tissue. To enhance cell lysis, two freeze-thaw steps were performed by freezing samples in a −20 C freezer for 2 h, and thawing at 60 C for 1 h. Then, 250 μL of 1.25 N HCl were added to each sample, and incubated at 60 C for 15 min. A 25 μL aliquot was transferred to microtiter plates, and 100 μL of 0.25% (w/v) periodic acid and 0.25% (w/v) sodium *m*-periodate was added and samples were incubated at room temperature for 90 min. To stop shikimate oxidation, 100 μL of 0.6 N NaOH and 0.22 M sodium sulfite were added. Shikimate was quantified at 380 nm using a spectrophotometer, and data were analyzed by fitting a standard curve of technical grade shikimate and subtracting background absorbance from samples. Data are presented in ng shikimate μg^-1^ fresh weight (FW). Survival data were also collected 30 days after glyphosate treatment by giving a “0” for plants that survived, and “1” to individuals that died.

### High Throughput Sequencing

A genotype-by-sequencing study was performed to identify SNPs for population genetic analyses. DNA was extracted with a commercial kit following the manufacturer’s recommendations (Mag-Bind^®^ Plant DNA DS, Omega Bio-Tek, Norcross, GA, USA), followed by sample preparation according to method developed by Elshire et al. (2011). Briefly, 200 ng of DNA from each sample was digested with 10 U *ApeKI* (New England BioLabs Inc., Ipswich, MA, USA) for 2 h at 75 C. Barcodes (4-8 bp) and common adapters were ligated with 400 U *T4 DNA ligase* (New England Biolabs Inc.) at 22 C for 1 h, followed by enzyme inactivation at 65 C for 30 min. Samples were then multiplexed (96 samples per pool, total of three pools), and purified using a commercial kit (QiAquick PCR purification kit, QiaGen, Germantown, MD, USA). A PCR amplification step was performed using P1 and P2 as primers with 14 cycles (98C for 30 s, 14 cycles of 98 C for 10 s, 68 C for 30 s, and 72 C for 30 s), and a final extension at 72 C for 5 min (Phusion High-Fidelity PCR Master Mix, Thermo Scientific). A final library clean-up step was performed before sequencing (QiAquick PCR purification kit, QiaGen). Library quality control was performed with qPCR and a bioanalyzer (High Sensitivity DNA Analysis, Agilent, Santa Clara, CA, USA). Sequencing (three libraries of 96 multiplexed samples each) was performed with a HiSeq3000 in 150 bp paired-end mode at the Center for Genome Research and Biocomputing (CGRB) at Oregon State University.

High-throughput sequencing data were processed with Stacks 2.55 (Rochette et al., 2019). Samples were demultiplexed with the *process_radtags* module, with the *--paired, -c, -q*, and *-r* flags. *De novo* assembly of loci was optimized as recommended by Paris et al. (2017). First, forward reads were assembled *de novo* with the *ustacks* module (*M*=4, *m*=3, *N*=2). Second, *cstacks* was used to build catalog loci. Third, *sstacks* was used to align the *de novo* loci to the catalog. Fourth, *tsv2bam* was implemented to transpose sequencing data to be oriented by locus, and paired-end reads were integrated to each single-end locus assembled. Finally, SNPs were called with the *gstacks* module. An integrated approach was used to align the catalog loci to the draft genome of *Lolium perenne*, a close relative to *L. multiflorum* (Byrne et al., 2015). We integrated the alignments back into the *gstacks* files using the *stacks-integrate-alignments* program and obtained genomic coordinates. Building *de novo* loci, followed by alignment of consensus sequences to the draft genome of *L. perenne*, combines the advantages of the *de novo* approach with the positional data from the draft genome (Paris et al., 2017).

SNPs were initially filtered with the *populations* module of Stacks (*--min-maf* 0.05, --*max-obs-het* 0.7, and *--hwe*), followed by *vcftools* (Danecek et al., 2011) to exclude sites that were missing in more than 90% of the individuals, and individuals with more than 90% missing sites. This dataset was used for the population genetics analyses described below.

### Sanger sequencing

Amino acid substitutions in EPSPS at position 102 and/or 106 have been demonstrated to cause conformational changes in the glyphosate target enzyme, preventing inhibition (reviewed by (Sammons & Gaines, 2014). We used Sanger sequencing to test whether these previously identified mutations in the *EPSPS* gene also played a role in glyphosate resistance in the Oregon populations of *L. multiflorum*. DNA from the same samples was used to amplify a 338 bp fragment of the *EPSPS* gene containing positions 102 and 106, which is located in exon 2 of this 1536-bp long gene (based on the coding sequence of *Oryza sativa*). Five to six individuals were sequenced per population. We used primers described by Adu-Yeboah et al. (2014) for PCR amplification (Platinum *Taq* DNA Polymerase High Fidelity, Invitrogen) following the manufacturer’s recommendations. BigDye Terminator v3.1 (Applied Biosystems, Bervely, MA, USA) was used for sequencing in an ABI 3730 sequencer (Applied Biosystems Inc.). To determine if resistant individuals were differentiated from susceptible individuals, we performed a multiple alignment in Geneious Prime 2020.0.4 (www.geneious.com) with a reference *EPSPS* sequence from *L. multiflorum* (Perez-Jones et al., 2007).

### Copy number variation and gene expression analysis

Copy number variation of *EPSPS* has been identified to confer glyphosate resistance in several weed populations, including *L. multiflorum* from Arkansas (Salas et al., 2012). More recently, an ABC-transporter was shown to be constitutively up-regulated in the weed *Echinochloa colona* (Pan et al., 2021). This transporter is localized to the cell plasma membranes and is believed to be involved in the efflux of glyphosate from the cytoplasm into the apoplast.

Primers were designed to amplify a 68-bp fragment in the coding region of *EPSPS* from the *L. multiflorum* populations from our study (Supporting Information Table S2), as well as a 135-bp fragment of the *ALS* that was chosen as a housekeeping gene (Brunharo et al., 2019; Dillon et al., 2017). Five individuals were analyzed from each of the resistant populations and two susceptible populations (“Gulf’ and “lm_105”). We used genomic DNA for the *EPSPS* copy number variation analysis. Reactions consisted of 5 μL of SsoAdvanced Universal SYBR^®^ Green Supermix (BioRad, Hercules, CA, USA), 0.25 μL of forward and 0.25 μL of reverse primer at 10 μM, and 2 μL of genomic DNA normalized to 5 ng μL^-1^, and were performed in a OneStepPlus™ q-RT-PCR system (Applied Biosystems Inc). Amplification was carried out with an initial denaturation cycle at 98 C for 3 min, and 40 cycles of 98 C for 15 s, followed by 64 C for 60 s. Melt curves were generated to assess specificity of primers, and reaction products were run in an agarose gel at 1% to assess fragment size and number. Primer efficiency was also performed with both primer sets. *EPSPS* copy number from resistant populations, as well as lm_105, were compared to Gulf.

For quantifying the gene expression of the ABC transporter gene, *ABCC8*, we designed several primer pairs to amplify a region of this gene based on the available sequence from *E. colona* (NCBI accession number MT249005.1). After Sanger sequencing a 420-bp fragment from *L. multiflorum*, we designed shorter, *L. multiflorum-specific* primers for quantitative real time-PCR (Supporting Information Table S2). *ALS* was also used as housekeeping gene to normalize the expression levels of *ABCC8*. Approximately 50 mg of leaf tissue was sampled from plants at the 3-leaf stage and immediately frozen in liquid nitrogen. Five individuals were analyzed from each of the resistant populations and two susceptible populations (“Gulf” and “lm_105”). RNA was extracted using a commercial kit (RNseasy Plant Mini Kit, Qiagen), followed by cDNA synthesis (iSCRIPT, Bio-Rad). The q-RT-PCR was performed as described for *EPSPS*. Population PRHC was not included in this analysis. There were five biological replicates for each population, and the experiment was performed three times. The experimental runs were pooled into one dataset based on a Levene’s test of homogeneity of variance (P = 0.64). Copy number variation and gene expression were quantified using the 2^-ΔΔCt^ method (Schmittgen & Livak, 2008) and multiple comparisons were performed using Tukey’s contrasts (*glht* function) in R with a Bonferroni correction was applied to the P-values considering 10 populations. *ABBC8* gene expression from resistant populations, as well as lm_105, were compared to Gulf.

### Population Differentiation

Principal component analysis (PCA) was implemented to obtain an overview of the population structure among the *L. multiflorum* populations. PCA is a model-free data summary technique, enabling the identification of population structure regardless of the historical underlying process shaping present levels of genetic variability (McVean, 2009). Separate analyses were performed that included (a) the entire dataset, containing Oregon populations, Gulf, the California population, and perennial, and (b) Oregon populations and Gulf only. The *prcomp* function was applied to a scaled SNP dataset using a custom R script, and eigenvalues were plotted with ggplot2 (Wickham, 2016).

To further dissect the historical demographic events in the study populations, we inferred patterns of ancestry and admixture using ADMIXTURE 1.3.0 (Alexander et al., 2009). We again ran the analyses using two different datasets, similar to the PCA: the first consisting of all sampled populations (including Gulf, PRHC, and perennial), and the second without PRHC or the perennial. Prior to analysis, the *vcf* was sorted and converted to Hapmap format using TASSEL (Bradbury et al., 2007), before further conversion to a binary format with PLINK (Purcell et al., 2007). ADMIXTURE was run with multiple values of *K* (1-10) as outlined in Liu et al. (2020), and the *Q* scores reflecting the probability of assignment of each individual to cluster *K* from the two analyses were plotted with PONG (Behr et al., 2016). Under a scenario where resistance evolved once from a single common ancestor and then spread throughout the region, all resistant populations should show similar patterns of ancestry. However, if resistance evolved on multiple genetic backgrounds independently, then resistant populations should show little grouping at different levels of *K*. An alternative explanation is that recent gene flow between populations has led to highly admixed patterns of ancestry among resistant populations.

To further reveal whether different resistant populations shared a recent common ancestor, we used a distance-based phylogenetic analysis using pairwise F_ST_ values among all population pairs produced from Stacks (Catchen et al., 2013). To determine if resistant populations grouped together, as would be expected if there was a single origin of resistant individuals, the Neighbor function (Felsenstein, 2005) from *Phylip* was used to produce a population tree, and heatmaps of pairwise F_ST_ were constructed.

In an attempt to identify genomic regions under selection, we performed an F_ST_ analysis between all pairs of resistant and susceptible populations. Given the clear adaptive benefit of glyphosate resistance, natural selection at resistance loci should result in locally elevated genetic divergence between resistant and susceptive populations. Moreover, if resistance was conferred by a single locus that was shared among all resistant populations, then the same highly differentiated locus should be observed in comparisons between all resistant and susceptible populations. Conversely, if multiple F_ST_ outliers are observed, then this suggests a more complex genetic architecture is involved, with potentially distinct resistance mechanisms and independent origins.

Genome-wide pairwise F_ST_ values were obtained with the *--fstats* flag of the *populations* module in Stacks. Missing genotypes were imputed with LinkImputeR (Money et al., 2017) to increase the number of loci analyzed (accuracy = 0.93, correlation = 0.8). A similar approach to impute missing genotypes prior to identifying loci involved in local adaptation was adopted elsewhere (Pritchard et al., 2018). For all possible comparisons between each resistant and susceptible population, we extracted the top 1% of the F_ST_ distribution and searched for loci that were found to overlap among multiple comparisons.

### Outlier annotation

The previous analysis aimed to identify loci under selection that could harbor glyphosate resistance genes. To determine potential loci involved in glyphosate resistance, we annotated the genomic contigs containing these outlier loci. For all loci that were found in common in the top 1% of most differentiated SNPs between a single resistant population and each of the susceptible populations, we extracted the entire contig containing that site from the *L. perenne* draft genome (Byrne et al., 2015). Augustus was used to predict genes within these contigs using an *Arabidopsis thaliana* trained dataset (Stanke et al., 2006). Predicted genes were annotated with Blast2GO 5 (Götz et al., 2008) using the *nr* database from NCBI, with an E-value cutoff of 10^-10^.

## RESULTS

### Glyphosate resistance is widespread

Our approach to phenotype *L. multiflorum* populations provided a clear distinction between resistant and susceptible individuals, because the 1456 g e.a. ha^-1^ glyphosate dose killed all individuals from the known susceptible population (Gulf), whereas 100% survival was observed for the known glyphosate-resistant population (PRHC). Out of the 16 Oregon field populations analyzed, eight were glyphosate resistant and were characterized by an exceptionally low accumulation of shikimate (Figure 2). Susceptibility or resistance to glyphosate based on shikimate accumulation was largely uniform among individuals within populations (Supporting Information Table S1). Susceptible individuals consistently accumulated >50 μg g^-1^ FW of shikimate, whereas resistant individuals accumulated <10 μg g^-1^ FW. The qualitative nature of shikimate accumulation between resistant and susceptible populations suggests there is a simple genetic basis for glyphosate resistance. This has been observed in *L. multiflorum* populations from New Zealand exhibiting non-target-site resistance (Ghanizadeh & Harrington, 2018). The shikimate accumulation data were highly consistent with data on survival, with mortality almost always occurring in individuals that accumulated high levels of shikimate. For a single individual in populations lm_24 and lm_60, and two each from lm_48 and lm_53, shikimate and survival data were not consistent. These individuals were excluded from further analysis.

**Fig. 2.**
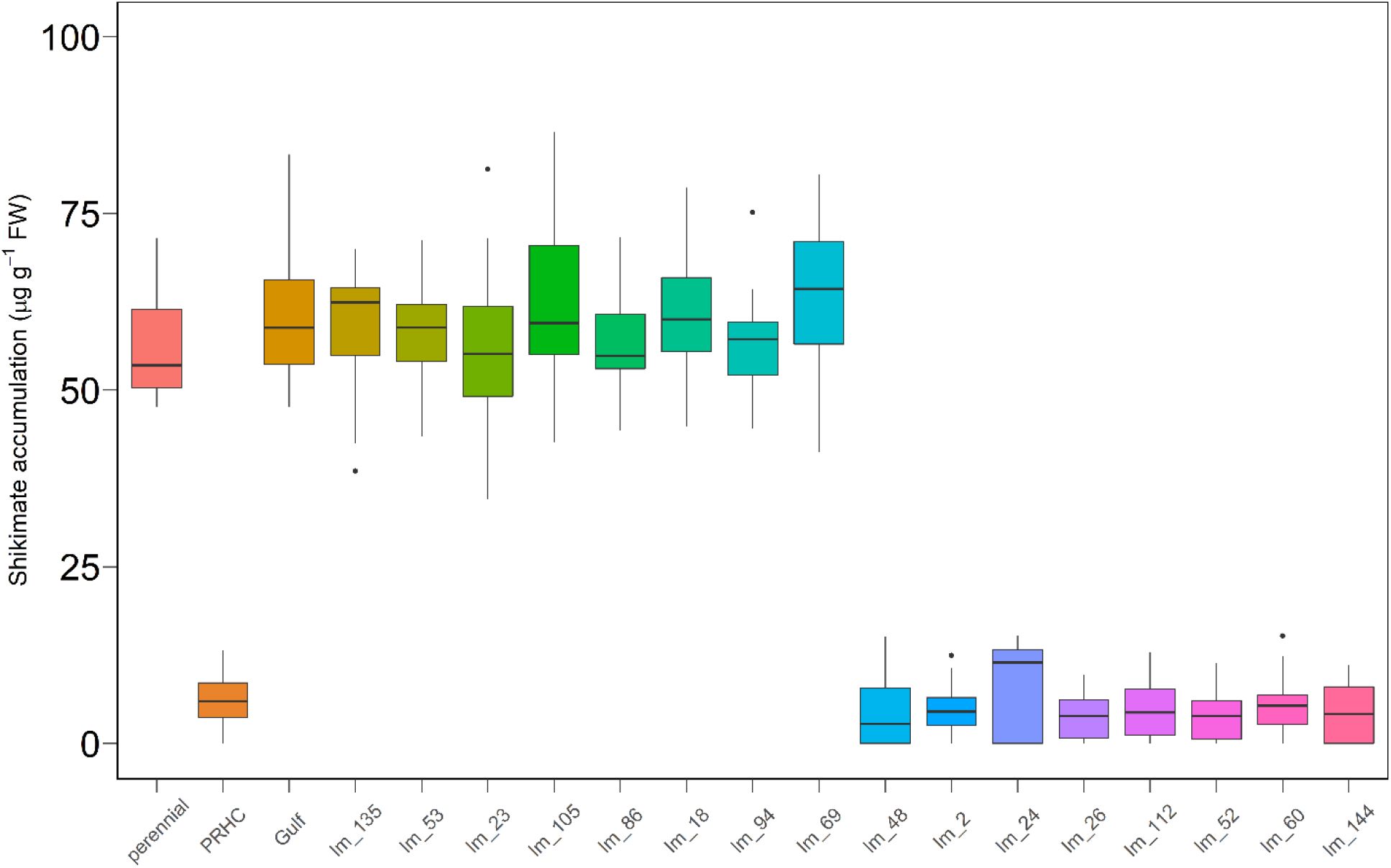
Shikimate accumulation in glyphosate resistant and susceptible *L. multiflorum* 48 hours after glyphosate treatment at 1456 g e.a. ha^-1^. Horizontal lines correspond to the median accumulation, box heights indicate the lower and upper quartile, and whiskers correspond to 1.5 times the interquartile range.

### No evidence for known resistance mechanisms in Oregon populations of *L. multiflorum*

The sequence analysis of *EPSPS* in Oregon populations of *L. multiflorum* revealed no evidence of mutations at positions 102 or 106, which were shown previously to be associated with the resistance phenotype (NCBI accession numbers MZ418136 and MZ418137). PRHC was included as a positive control for the presence of functionally-relevant amino acid substitutions in EPSPS at position 106 that confer resistance. As expected, a mutation was found in *EPSPS* at position 106 from PRHC, causing a proline-to-alanine substitution. There were synonymous mutations found among the sequenced individuals from Oregon, but no mutation was found in common among all resistant individuals sequenced and no non-synonymous mutations were observed.

Little variation was observed in the number of *EPSPS* copies across the surveyed populations relative to the housekeeping gene *ALS* (Figure 3). After normalizing to *ALS*, mean and median *EPSPS* copy numbers across Oregon populations relative to Gulf were 1.13 and 0.99, respectively, and no statistically significant differences were observed. Moreover, we did not observe differences between resistant and susceptible individuals in the expression levels of the ABC transporter *ABCC8* in the Oregon *L. multiflorum* populations (Figure 4), with mean and median *ABCC8* expression of 0.93 and 0.81 relative to Gulf. These results suggest a novel glyphosate resistance mechanism exists in *L. multiflorum* populations from Oregon.

**Fig. 3.**
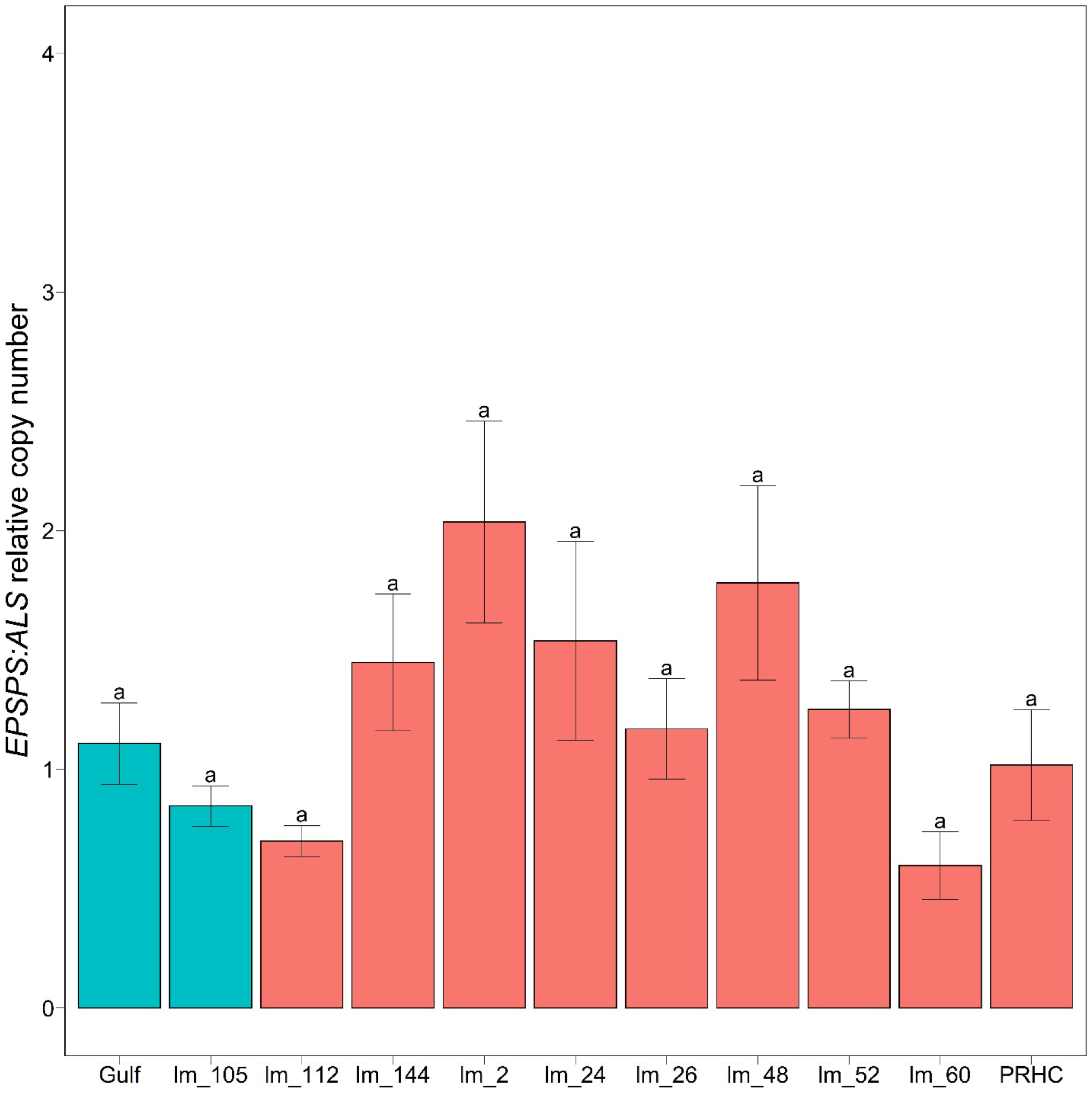
Copy number variation of *EPSPS* among *L. multiflorum* populations relative to *ALS*. Bars represent standard errors around the mean. Copy number variation was quantified using the 2^-ΔΔCt^ method. No significant difference was detected between resistant and susceptible populations. Multiple comparisons were performed using Tukey’s contrasts, with Bonferroni-corrected p-values considering an α = 0.05. Green bars indicate susceptible populations, and red bars correspond to resistant populations.

**Fig. 4.**
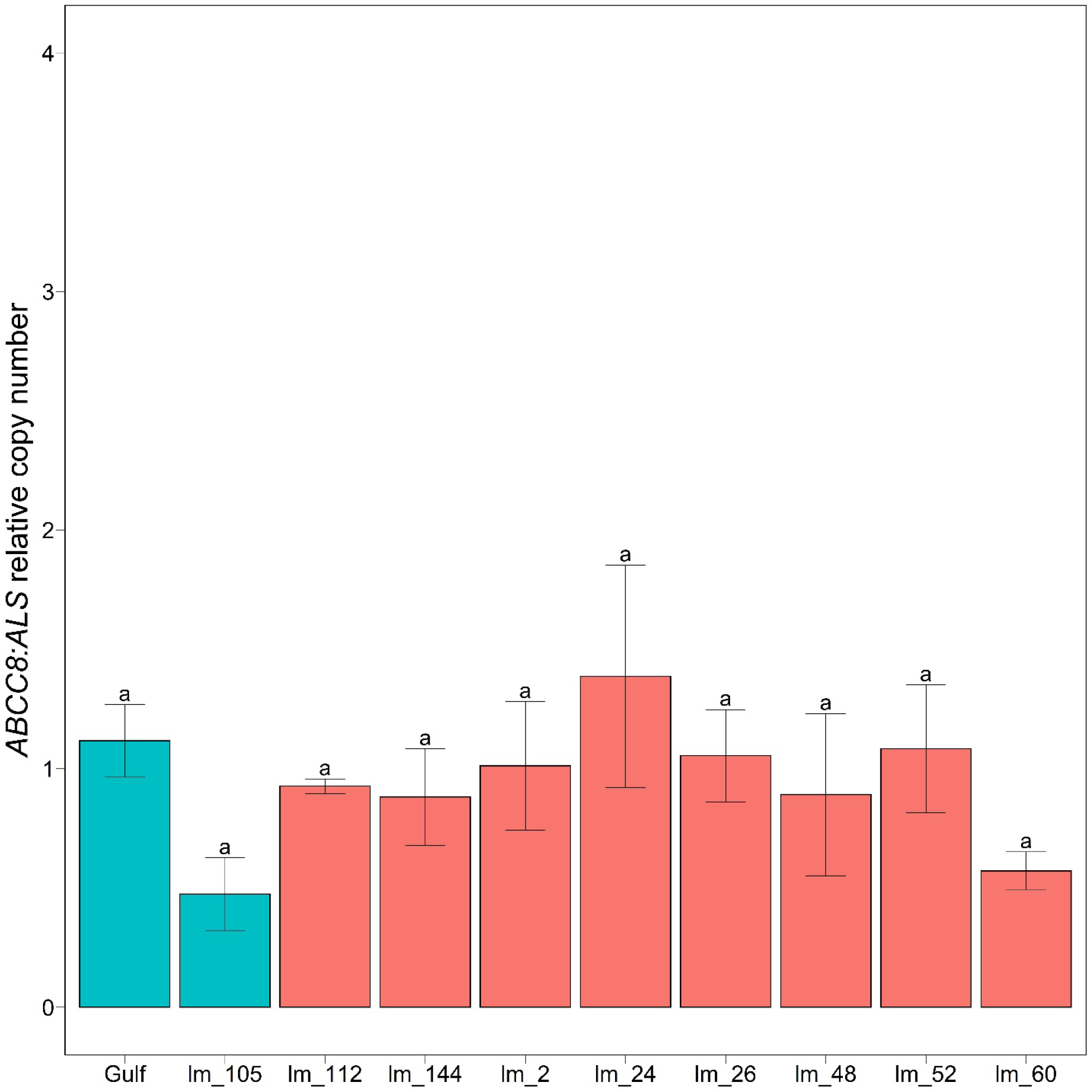
Expression of the ATP-binding cassette *ABCC8* in two glyphosate-susceptible populations (green) and eight –resistant (red) of *L. multiflorum* relative to the housekeeping gene *ALS*. Bars represent standard errors around the mean. Gene expression variation was quantified using the 2^-ΔΔCt^ method. No significant difference was detected between resistant and susceptible populations. Multiple comparisons were performed using Tukey’s contrasts, with Bonferroni-corrected p-values considering an α = 0.05. Green bars indicate susceptible populations, and red bars correspond to resistant populations.

### Little genetic structuring of resistant and susceptible populations

After removing reads with adaptor sequence, without barcodes and restriction enzyme cut sites, and discarding low quality reads, the *process_radtags* program retained 407M (SE = 6.7M) reads that were used for the *Stacks* pipeline. Following the filtering steps in the *populations* module and vcftools, 34,933 SNP’s were retained for population genetics analyses.

Patterns of genetic variation among individuals and populations are structured primarily according to geography and species, rather than their resistance phenotype. In the analysis of the entire dataset, the first principal component mainly reveals differentiation between the two different species: the annual and perennial ryegrass. Similarly, PC2 largely separates the geographically distant PRHC population from the Oregon populations. Among the Oregon populations only, PCA explains little of the genetic variation present (the first two principal components explain a combined 3% of the variation) (Figure 5B). Moreover, there is little separation between the resistant and susceptible individuals along the first two principal components, suggesting a recent common ancestor of these individuals and/or ongoing gene flow between them. Consistent with the PCA, average pairwise F_ST_ indicates little differentiation among populations in Oregon (mean = 0.09, median = 0.09) (Figure S1).

**Fig. 5.**
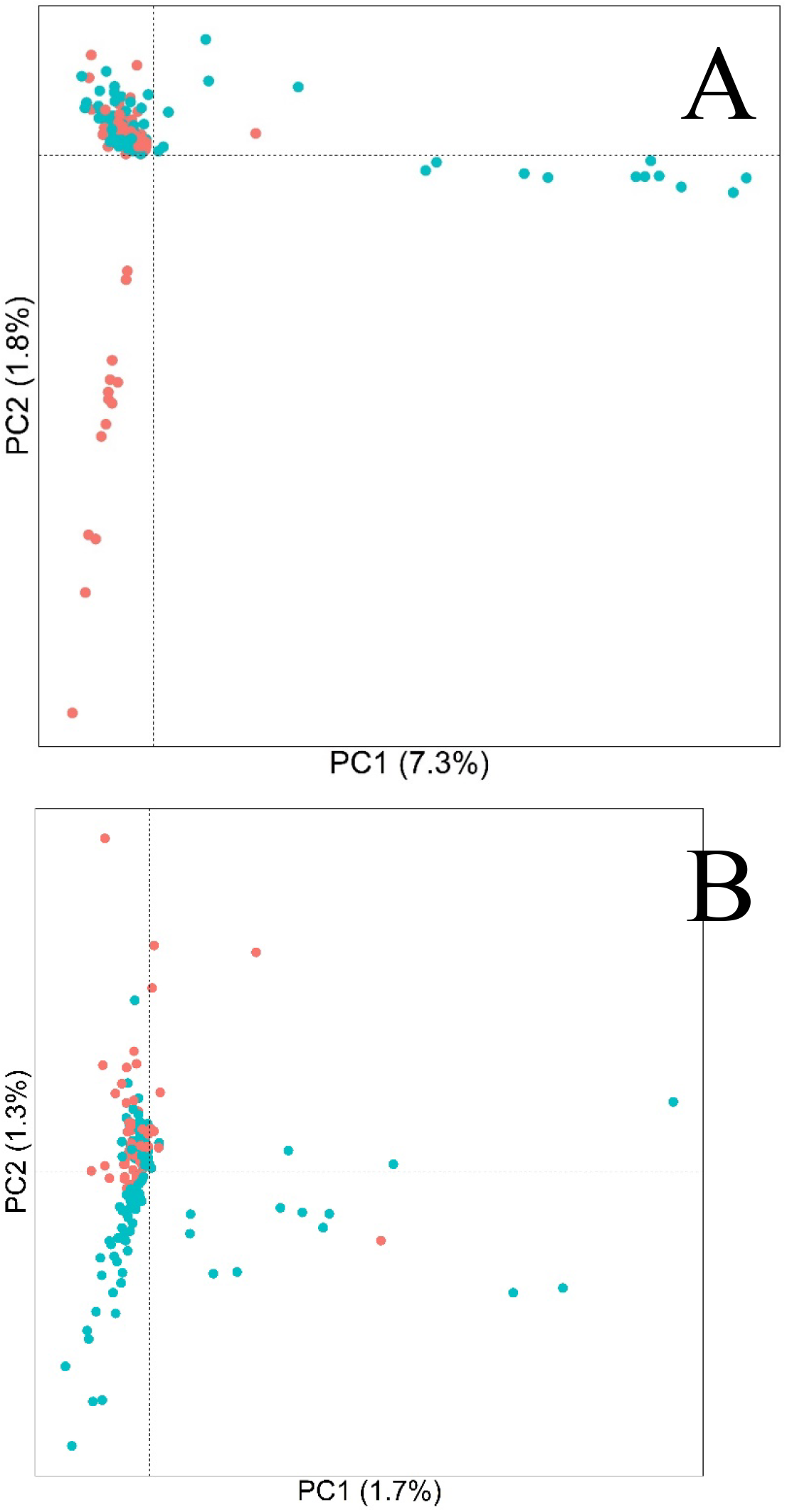
Principal Component Analysis (PCA) from 34933 SNPs among individuals from resistant (red) and susceptible (green) ryegrass weed populations. (A) Analysis with all populations, including the 16 *L. multiflorum* populations from Oregon and one population from California (PRHC), as well as a single population of the perennial ryegrass *L. perenne*. (B) Analysis of only the Oregon *L. multiflorum* populations.

The results from ADMIXTURE are largely consistent with the PCA. The full dataset indicates that *L. multiflorum* populations share little ancestry with the perennial species (*L. perenne*) (Figure 6A). However, some populations do exhibit shared ancestry with the perennial species (particularly population lm_69). This can be explained by hybridization and introgression between these species, as both species co-exist and remain inter-fertile. At higher values of *K*, clear differentiation is found between the Oregon populations and the California *L. multiflorum* population (PRHC). When *L. perenne* and PRHC are removed from the analysis, additional differences are observed among the Oregon populations. At *K=2*, the resistant and susceptible individuals are not assigned to separate clusters. Rather, consistent with the PCA, there is extensive shared ancestry among these populations, further supporting a recent common ancestor among all of the Oregon populations. However, at *K* = 6, the glyphosate-resistant populations form multiple distinct groups. Despite some shared ancestry between the resistant and susceptible populations, susceptible populations do not show similar levels of population structure. In addition, there is little evidence of admixture among resistant populations, with only six of the resistant individuals showing any evidence of admixture. These results suggest that ongoing gene flow is unlikely to explain differences in ancestry among the resistant populations, and that glyphosate resistance likely has evolved independently on different genetic backgrounds.

**Fig. 6.**
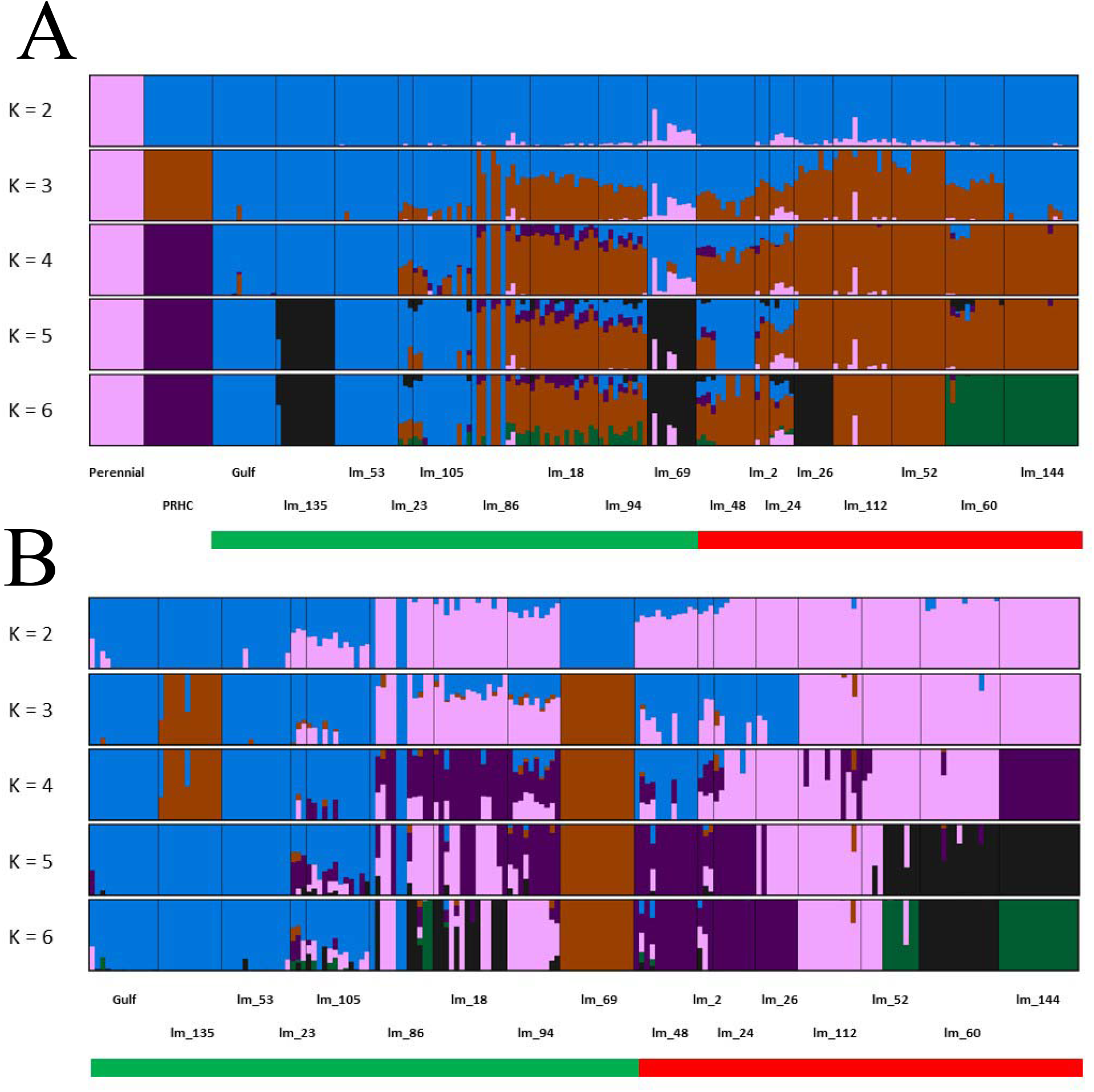
Ancestry coefficients from *Admixture* showing assignment probabilities into *K* = 2 to 6 different clusters. Analysis with the full dataset (A), and without *L. perenne* and California populations (PRHC) (B). The green bar represents glyphosate-susceptible populations, whereas red represents glyphosate-resistant.

Patterns of genetic relatedness among populations based on pairwise genetic distances also reveal little evidence of structuring between resistant and susceptible populations. Rather, resistant populations are inter-digitated with susceptible populations, further suggesting multiple origins of the resistant phenotype (Figure S2). Thus, all three analyses provide consistent findings of little overall population structure among Oregon populations of *L. multiflorum*, but the presence of divergence among resistant populations suggests that resistance evolved on multiple distinct genetic backgrounds.

### Patterns of genetic differentiation do not support a single origin of resistance

Between each pair of resistant and susceptible populations, overall levels of population differentiation (F_ST_) were quite low. Median F_ST_ varied from 0.022 to 0.032, depending on the population, with most of the third quartile of the distribution below 0.067 (Figure 7). Despite the overall low median F_ST_ among the glyphosate-resistant populations analyzed, on average, 0.86% of the loci revealed F_ST_ above 0.5, with one SNP in locus 23735 reaching F_ST_ of 1 in population lm_2. Overall, there were no loci that were consistently found in the top 1% of the F_ST_ distribution for all resistant and susceptible population pairs. Therefore, we next asked whether there were any shared outliers in comparisons between a single resistant population and each of the susceptible populations. Resistant populations lm_2, lm_24, lm_26, lm_60, lm_112, and lm_144 exhibited 16, 2, 2, 4, 1 and 1 loci, respectively, that were ranked in the top 1% of the distribution for all pairwise comparisons. Populations lm_48, lm_52, and lm_60 did not have any SNPs that were consistently ranked in the top 1% of the F_ST_ distribution that were shared among all susceptible populations. Only population lm_2 exhibited F_ST_ of 1 (Figure 7). The predicted proteins encoded by genes found on the contigs containing these loci included transmembrane transporters, protein kinase (L-type lectin-domain containing receptor kinase IX.2-like), anion transporters (GABA transporter 1), and enzymes previously reported to be able to metabolize herbicides (P450s), among other hypothetical proteins of unknown function (Supporting Information Table S4).

**Fig. 7.**
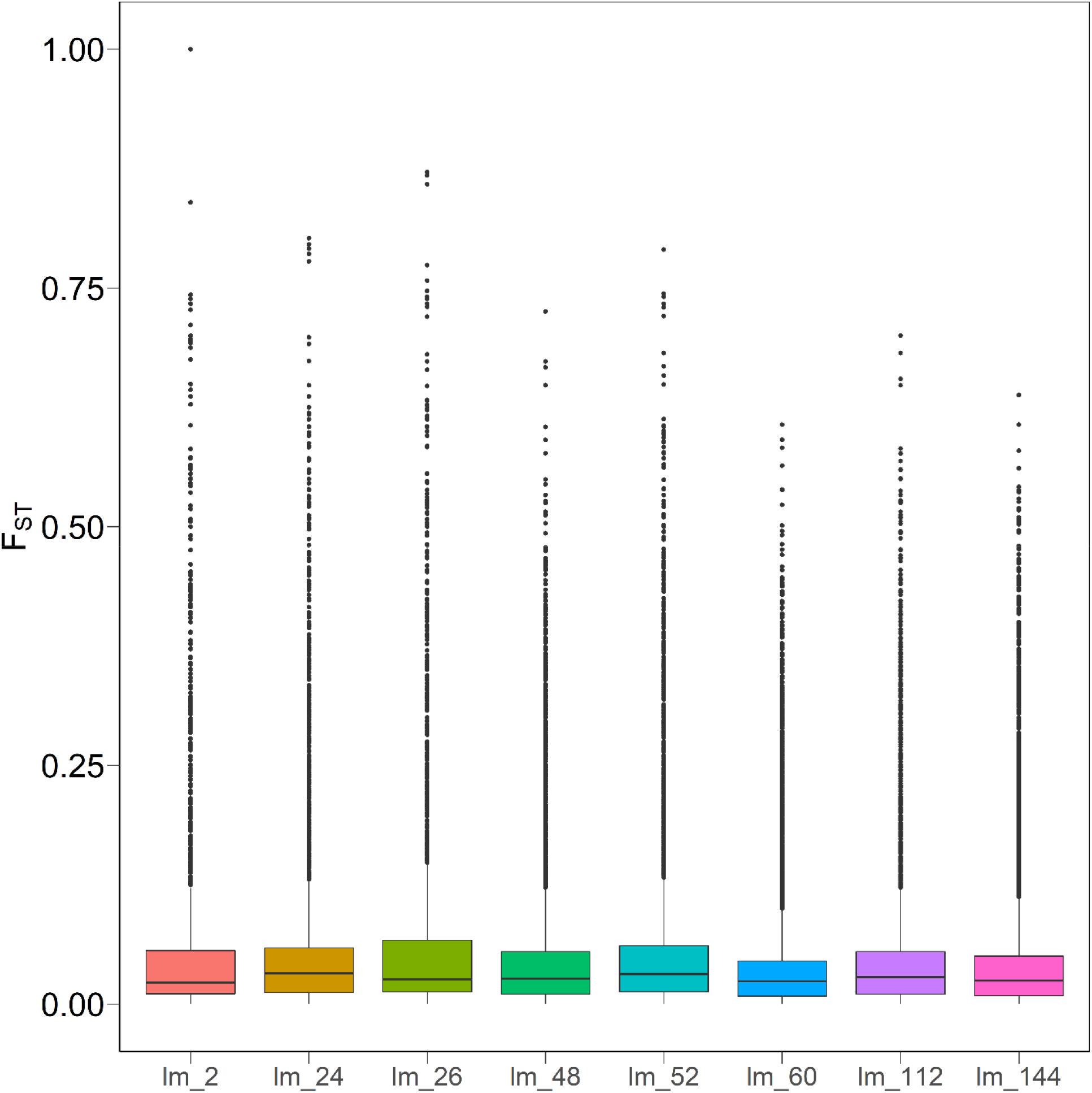
Distribution of F_ST_ values for all SNPs between each glyphosate-resistant *L. multiflorum* population (listed) and each of the susceptible populations. Horizontal lines correspond to the median F_ST_, box heights indicate the lower and upper quartile, and whiskers correspond to 1.5 times the interquartile range. Outliers outside this range are depicted as points.

## DISCUSSION

The evolution of herbicide resistance is a growing challenge to broad acreage agriculture that depend on herbicides for weed management. In many cases, alternative control methods are costly or unavailable, and chemical companies have not introduced many new herbicides over the past few decades. Herbicide resistance in weeds is a clear example of rapid adaptation caused by repeated, strong selection pressures induced by human intervention. Our findings reveal multiple instances of the evolution of glyphosate resistance in *L. multiflorum* across different fields in Oregon. Despite frequent examples of resistance evolving due to mutations that impact the target site, our sequence and population genomic analyses imply a more complex history of resistance.

Amino acid substitutions at positions 102 and 106 in EPSPS have been shown to confer resistance to glyphosate in *L. multiflorum* (Brunharo & Hanson, 2018) and *E. colona* (Morran et al., 2018) in California, *Chloris virgata* in Australia (Ngo et al., 2017), *Amaranthus tuberculatus* from Mississippi (Nandula et al., 2013), and others. However, in the resistant Oregon *L. multiflorum* populations we sequenced, no missense mutations in *EPSPS* were found. Moreover, our results suggest that copy number variation in *EPSPS* is not involved in glyphosate resistance (Figure 3). Increased copy number leads to increased dosage of EPSPS protein in plant cells, requiring higher concentrations of glyphosate to inhibit the enzyme (Powles, 2010). Copy number differences in *EPSPS* that confer glyphosate resistance have occurred repeatedly in several weed species, likely due to different molecular mechanisms. For example, in *Amaranthus palmeri*, resistant plants had up to 160-fold more copies of *EPSPS* than susceptible individuals, likely due to extrachromosomal circular DNA (Koo et al., 2018). In addition, resistant *Kochia scoparia* plants had up to 10 copies of the gene (Gaines et al., 2016), likely mediated by a mobile genetic element (Patterson et al., 2019). Lastly, *EPSPS* duplication associated with a nonsense mutation in the *EPSPS* gene has been observed in the allotetraploid grass species *Poa annua* (Brunharo et al., 2018).

A plasma membrane-localized ABC transporter has been recently identified to confer glyphosate resistance in an *E. colona* population from Australia (Pan et al., 2021). Data suggest that this transporter enhances glyphosate efflux from the cytoplasm into the apoplast. Further functional characterization was confirmed by transforming several plants to over-express the *ABCC8* gene, which conferred tolerance to field rates of the herbicide. Our results show no evidence of differences in *ABCC8* expression between resistant and susceptible populations of *L. multiflorum* in Oregon. Thus, despite the widespread involvement of genetic changes in *EPSPS* resulting in glyphosate target-site resistance, a different mechanism, likely involving a non-target-site basis, appears to confer resistance in Oregon populations of *L. multiflorum*.

There are at least three different processes that could explain the presence of multiple glyphosate resistant *L. multiflorum* populations found in different fields across Oregon. First, there could be a single origin of resistance, followed by human-assisted dispersal of the resistance phenotype. Alternatively, there could be a single origin, followed by gene flow among neighboring populations. Finally, there could be multiple origins of resistance due to independent mutations in the same or different genes. Our results favor the third explanation. If there was a single origin of resistance, then we would expect all resistant individuals to group together in our analyses. However, the PCA showed no clustering of resistant and susceptible individuals. Similarly, at *K*=2, there was no evidence of distinct ancestry patterns associated with the resistance phenotype. Finally, the tree-based analysis showed resistant and susceptible populations interspersed along the tree. Similarly, we can rule out a single origin followed by gene flow among populations, because *L. multiflorum* was introduced to Oregon in the early 1900s, and glyphosate did not become adopted as a widespread herbicide until the year 2000 (Benbrook, 2016). Therefore, it is unlikely that there would have been sufficient time for glyphosate-resistance gene(s) to spread naturally via gene flow across this expansive region. By contrast, at higher values of *K*, distinct patterns of ancestry emerged in each resistant population but not in the susceptible populations. Given that these populations are all resistant to the same herbicide but have different patterns of ancestry provides strong evidence that resistance evolved separately on different genetic backgrounds. Confirming this hypothesis will require future characterization of the gene or genes that confer resistance in these populations.

Consistent with a multiple-origins model, our analysis of genetic divergence across the genome of *L. multiflorum* fails to find loci that were repeatedly differentiated between resistant and susceptible populations. Under a model of a single origin of resistance and repeated, strong, and recent selection, we would expect to find variants at high frequency in the resistant individuals that are at low frequency in the susceptible plants. This should be manifest as high F_ST_ at SNPs linked to the mutation conferring resistance. Moreover, this locus should show consistently high F_ST_ between each pair of resistant and susceptible populations. However, we found no SNPs in our dataset that were repeatedly at high F_ST_ between different pairs of resistant and susceptible populations. Although the obligate outcrossing, annual life history of *L. multiflorum* and the reduced representation genotyping approach used here limit the genomic resolution and the extent of linkage disequilibrium (LD) in the dataset, strong and recent natural selection should result in long blocks of LD between SNPs reasonably tightly linked to functional mutations. By contrast, it is likely that the diverse patterns seen in these resistant Oregon populations are due to mutations (maybe at different loci) that pre-dated the widespread use of glyphosate and segregated at low frequency in the ancestral population. To test this scenario, a different approach aimed at detecting selection on standing genetic variation would be better suited than one that searches for shared outliers. However, the lack of shared outliers is consistent with our analyses of population structure, further supporting the conclusion that resistance does not have a single origin in these Oregon populations. Given the apparent complexity of the origin of resistance in these populations, future studies that sequence whole genomes from these populations will be necessary to test these alternate hypotheses.

Even though no outliers were consistently found between all pairs of resistant and susceptible individuals, we did find SNPs with high F_ST_ that were in common between a single resistant population and each of the susceptible populations. Gene annotation and ontology analyses of genomic contigs containing these genes identified several molecular functions potentially associated with resistance. For example, we detected genes involved in the detoxification of xenobiotics, most notably cytochrome P450 genes. Genes in this family have been suggested to be involved in glyphosate resistance by enhancing herbicide degradation in other populations of *L. multiflorum, L. rigidum*, and *L. perenne* outside of Oregon (Suzukawa et al., 2021). In addition, we identified genes involved in transmembrane transport (Supporting Information Table S4). Vacuolar sequestration of glyphosate has been suggested to confer resistance in other species (Peng et al., 2010). Although much research has been conducted to elucidate non-target-site resistance mechanisms to glyphosate in *L. multiflorum* and other weed species, the genetic and molecular mechanisms remain largely unknown (Suzukawa et al., 2021).

This research suggests that glyphosate resistance evolved multiple times in *L. multiflorum* populations from Oregon. Elucidating the molecular mechanisms of herbicide resistance is crucial for the improvement of weed management practices. Although target-site resistance has been described frequently for many herbicides with different mechanisms of action, the underlying molecular characteristics that confer non-target-site resistance remain largely unknown (Baucom, 2016; Suzukawa et al., 2021). Identification of the genetic changes involved in resistance evolution will allow the development of quick herbicide resistance diagnostics in the laboratory and in the field. The rapid diagnosis of herbicide resistance will allow the development of measures that slow down, or preferably prevent, the introduction of resistance alleles to new areas, reducing the long-term costs associated with herbicide resistance. Finally, because non-target-site resistance may confer resistance to herbicides from different chemical groups (i.e. generalist resistance mechanisms; Comont et al., 2020), a more in-depth understanding of these mechanisms will allow better utilization of herbicides to manage the spread of resistance.

## ACKNOWLEDGEMENTS

CAB would like to thank the Oregon Department of Agriculture, Oregon Seed Council, and Oregon State University Agricultural Research Foundation for partial funding provided for this research.

## AUTHOR CONTRIBUTION

CAB designed the research and performed the experiments. CAB and MAS performed the statistical analysis, data interpretation, and manuscript preparation. CAB and MAS contributed equally.

## DATA AVAILABILITY

The genotype-by-sequencing data used in this study are available at NCBI SRA PRJNA739185. *EPSPS* and *ABCC8* sequences are available at NCBI (accession numbers MZ418136 and MZ418137). Raw data used for the *EPSPS* amplification and *ABCC8* expression are available at https://github.com/caiobrunharo/popgen_ryegrass.

## Supporting Information

**Fig. S1**. Heatmap of pairwise F_ST_ among glyphosate-resistant and –susceptible *L. multiflorum* populations. Annual ryegrass variety Gulf was included in the analysis. Histogram with distribution of pairwise F_ST_ values from heatmap (mean = 0.09, median=0.09).

**Fig. S2.** Neighbor joining tree based on pairwise genetic distances (F_ST_) between all pairs of *L. multiflorum* populations from Oregon, using the perennial *L. perenne* as an outgroup. Red and blue are glyphosate-resistant (R) and –susceptible (S), respectively.

**Table S1.** Geographical location, resistance phenotype, sample size, and mean accumulated shikimate (± SE) for the populations in this study.

**Table S2.** Primers designed for copy number variation in *L. multiflorum* populations

**Table S3.** Loci within the top 1% of the F_ST_ distribution in all pairwise comparisons.

**Table S4.** Annotations of contigs with SNP’s exhibiting F_ST_ of 1. Contigs were subjected to AUGUSTUS analysis to predict coding proteins, and coding proteins were annotated with Blast2GO 5.

## REFERENCES

Alexander DH, Novembre J, Lange K. 2009. Fast model-based estimation of ancestry in unrelated individuals. Genome Research 19: 1655–1664.

Appleby AP, Olson PD, Colbert DR. 1976. Winter wheat yield reduction from interference by Italian ryegrass. Agronomy Journal 68: 463–466.

Baucom RS. 2016. The remarkable repeated evolution of herbicide resistance. American Journal of Botany 103: 181–183.

Baucom RS. 2019. Evolutionary and ecological insights from herbicide-resistant weeds: what have we learned about plant adaptation, and what is left to uncover? New Phytologist 223: 68–82.

Behr AA, Liu KZ, Liu-Fang G, Nakka P, Ramachandran S. 2016. pong: fast analysis and visualization of latent clusters in population genetic data. Bioinformatics 32: 2817–2823.

Bobadilla LK, Hulting AG, Berry PA, Moretti ML, Mallory-Smith C. 2021. Frequency, distribution, and ploidy diversity of herbicide-resistant Italian ryegrass *(Lolium perenne spp. multiflorum)* populations of western Oregon. Weed Science 69: 177–185.

Bolnick DI, Barrett RDH., Oke KB, Rennison DJ, Stuart YE. 2018. (Non)Parallel evolution. Annual Review of Ecology, Evolution, and Systematics 49: 303–330.

Bradbury PJ, Zhang Z, Kroon DE, Casstevens TM, Ramdoss Y, Buckler ES. 2007. TASSEL: software for association mapping of complex traits in diverse samples. Bioinformatics 23: 2633–2635.

Brunharo CACG, Hanson BD. 2018. Multiple herbicide-resistant Italian ryegrass *[Lolium perenne* L. *spp. multiflorum* (Lam.) Husnot] in California perennial crops: characterization, mechanism of resistance, and chemical management. Weed Science 66: 696–701.

Brunharo CACG, Morran S, Martin K, Moretti ML, Hanson BD. 2018. EPSPS duplication and mutation involved in glyphosate resistance in the allotetraploid weed species *Poa annua* L. Pest Management Science 75: 1663–1670.

Brunharo CACG, Takano HK, Mallory-Smith CA, Dayan FE, Hanson BD. 2019. *Role of glutamine synthetase isogenes and herbicide metabolism in the mechanism of resistance to glufosinate in *Lolium perenne* L. spp. multiflorum* biotypes from Oregon. Journal of Agricultural and Food Chemistry 67: 8431–8440.

Busi R, Powles SB. 2016. Cross-resistance to prosulfocarb + S-metolachlor and pyroxasulfone selected by either herbicide in Lolium rigidum. Pest Management Science 72: 1664–1672.

Byrne SL, Nagy I, Pfeifer M, Armstead I, Swain S, Studer B., Mayer K, Campbell JD, Czaban A, Hentrup S et al. 2015. A synteny-based draft genome sequence of the forage grass Lolium perenne. The Plant Journal 84: 816–826.

Catchen J, Bassham S, Wilson T, Currey M, O’Brien C, Yeates Q, Cresko WA. 2013. The population structure and recent colonization history of Oregon threespine stickleback determined using restriction-site associated DNA-sequencing. Molecular Ecology 22: 2864–2883.

Comont D, Lowe C, Hull R, Crook L, Hicks HL, Onkokesung N, Beffa R, Childs DZ, Edwards R., Freckleton RP et al. 2020. Evolution of generalist resistance to herbicide mixtures reveals a trade-off in resistance management. Nature Communications 11: e3086.

Danecek P, Auton A, Abecasis G, Albers CA, Banks E, DePristo MA, Handsaker RE, Lunter G, Marth GT, Sherry ST et al. (2011) The variant call format and VCFtools. Bioinformatics 27: 2156–2158.

Dayan FE, Owens DK, Corniani N, Silva FML, Watson SB, Howell J, Shaner DL. 2015. Biochemical markers and enzyme assays for herbicide mode of action and resistance studies. Weed Science 63: 23–63.

Delye C. 2013. Unravelling the genetic bases of non-target-site-based resistance (NTSR) to herbicides: a major challenge for weed science in the forthcoming decade. Pest Management Science 69: 176–187.

Delye C, Jasieniuk M, Le Corre V. 2013. Deciphering the evolution of herbicide resistance in weeds. Trends in Genetics 29: 649–658.

Dillon A, Varanasi VK, Danilova TV, Koo D-H, Nakka S, Peterson DE, Tranel PJ, Friebe B, Gill BS, Jugulam M. 2017. Physical mapping of amplified copies of the 5-enolpyruvylshikimate-3-phosphate synthase gene in glyphosate-resistant *Amaranthus tuberculatus*. Plant Physiology 173: 1226–1234.

Dimaano NG, Yamaguchi T, Fukunishi K, Tominaga T, Iwakami S. 2020. Functional characterization of cytochrome P450 CYP81A subfamily to disclose the pattern of cross-resistance in *Echinochloa phyllopogon*. Plant Molecular Biology 102: 403–416.

DiTomaso JM, Healy EA. 2007. Weeds of California and other western states. Oakland, CA: University of California Division of Agriculture and Natural Resources.

Dixon A, Comont D, Slavov GT, Neve P. Population genomics of selectively neutral genetic structure and herbicide resistance in UK populations of *Alopecurus myosuroides*. Pest Management Science 77: 1520–1529.

Elshire RJ, Glaubitz JC, Sun Q, Poland JA, Kawamoto K, Buckler ES, Mitchell SE. 2011. A robust, simple genotyping-by-sequencing (GBS) approach for high diversity species. PLoS ONE 6: e19379.

Endo M, Osakabe K, Ichikawa H, Toki S. 2006. Molecular characterization of true and ectopic gene targeting events at the acetolactate synthase gene in *Arabidopsis*. Plant and Cell Physiology 47: 372–379.

Felsenstein J. 2005. PHYLIP (Phylogeny Inference Package) (Version 3.6).

Funke T, Han H, Healy-Fried ML, Fischer M, Schonbrunn E. 2006. Molecular basis for the herbicide resistance of Roundup Ready crops. Proceedings of the National Academy of Sciences, 103(35), 13010–13015.

Gaines TA, Barker AL, Patterson EL, Westra P, Westra EP, Wilson RG, Jah P, Kumar V, Kniss AR. 2016. EPSPS gene copy number and whole-plant glyphosate resistance level in *Kochia scoparia*. PLOS ONE, 11(12), e0168295.

Gaines TA, Zhang W, Wang D, Bukun B, Chisholm ST, Shaner DL, Nissen SJ, Patzoldt WL, Tranel PJ, Culpepper AS et al. 2010. Gene amplification confers glyphosate resistance in *Amaranthus palmeri*.. Proceedings of the National Academy of Sciences of the United States of America 107: 1029–1034.

Götz S, García-Gómez JM, Terol J, Williams TD, Nagaraj SH, Nueda MJ, Robles M, Talon M, Dopazo J, Conesa A. 2008. High-throughput functional annotation and data mining with the Blast2GO suite. Nucleic Acids Research 36: 3420–3435.

Han H, Yu Q, Beffa R, González S, Maiwald F, Wang J, Powles SB. 2021. Cytochrome P450 CYP81A10v7 in *Lolium rigidum* confers metabolic resistance to herbicides across at least five modes of action. The Plant Journal 105: 79–92.

Heap I. 2021. The International Survey of Herbicide Resistant Weeds. http://www.weedscience.org. [accessed 18 June 2021]

Hess M, Barralis G, Bleiholder H, Buhr L, Eggers T, Hack H, Stauss R. 1997. Use of the extended BBCH scale - general for the descriptions of the growth stages of mono- and dicotyledonous weed species. Weed Research 37: 433–441.

Humphreys M, Feuerstein U, Vandewalle M, Baert J. 2010. Ryegrasses. In: Boller B, Posselt U, Veronesi F, eds. Fodder Crops and Amenity Grasses. New York: Springer, 211–260.

Koo D-H, Molin WT, Saski CA, Jiang J, Putta K, Jugulam M, Friebe B, Gill BS. 2018. Extrachromosomal circular DNA-based amplification and transmission of herbicide resistance in crop weed *Amaranthus palmeri*. Proceedings of the National Academy of Sciences 115: 3332–3337.

Kreiner JM, Stinchcombe JR, Wright SI. 2018. Population genomics of herbicide resistance: adaptation via evolutionary rescue. Annual Review of Plant Biology 69: 611–635.

Lee KM, Coop G. 2017. Distinguishing among modes of convergent adaptation using population genomic data. Genetics 207: 1591–1619.

Lee KM, Coop G. 2019. Population genomics perspectives on convergent adaptation. Philosophical Transactions of the Royal Society B-Biological Sciences 374: e20180236.

Liu C-C, Shringarpure S, Lange K, Novembre J. 2020. Exploring population structure with admixture models and principal component analysis. In: Dutheil JY, ed. Statistical Population Genomics. New York, NY: Springer, 67–86.

McVean G. 2009. A Genealogical interpretation of principal components analysis. PLoS Genetics 5: e1000686.

Money D, Migicovsky Z, Gardner K, Myles S. 2017. LinkImputeR: user-guided genotype calling and imputation for non-model organisms. BMC Genomics 18: e523.

Morran S, Moretti ML, Brunharo CA, Fischer AJ, Hanson BD. (2018). Multiple target site resistance to glyphosate in junglerice *(Echinochloa colona)* lines from California orchards. Pest Management Science 74: 2747–2753.

Nandula VK, Ray JD, Ribeiro DN, Pan Z, Reddy KN. 2013. Glyphosate resistance in tall waterhemp *(Amaranthus tuberculatus)* from Mississippi is due to both altered target-site and nontarget-site mechanisms. Weed Science 61: 374–383.

Ngo TD, Krishnan M, Boutsalis P, Gill G, Preston C. 2017. Target-site mutations conferring resistance to glyphosate in feathertop Rhodes grass *(Chloris virgata)* populations in Australia. Pest Management Science 74: 1094–1100.

Oerke EC. 2006. Crop losses to pests. Journal of Agricultural Science 144: 31–43.

Oliveira MC, Gaines TA, Dayan FE, Patterson EL, Jhala AJ, Knezevic SZ. 2018. Reversing resistance to tembotrione in an *Amaranthus tuberculatus* (var. rudis) population from Nebraska, USA with cytochrome P450 inhibitors. Pest Management Science 74: 2296–2305.

Paris JR, Stevens JR, Catchen JM. 2017. Lost in parameter space: a road map for stacks. Methods in Ecology and Evolution 8: 1360–1373.

Patterson EL, Saski CA, Sloan DB, Tranel PJ, Westra P, Gaines TA. 2019. The draft genome of *Kochia scoparia* and the mechanism of glyphosate resistance via transposon-mediated EPSPS tandem gene duplication. Genome Biology and Evolution 11: 2927–2940.

Patzoldt WL, Hager AG, McCormick JS, Tranel PJ. 2006. A codon deletion confers resistance to herbicides inhibiting protoporphyrinogen oxidase. Proceedings of the National Academy of Sciences of the United States of America 103: 12329–12334.

Peng Y, Abercrombie LL, Yuan JS, Riggins CW, Sammons RD, Tranel PJ, Stewart CN. 2010. Characterization of the horseweed *(Conyza canadensis)* transcriptome using GS-FLX 454 pyrosequencing and its application for expression analysis of candidate non-target herbicide resistance genes. Pest Management Science 66: 1053–1062.

Perez-Jones A, Park KW, Polge N, Colquhoun J, Mallory-Smith CA. 2007. Investigating the mechanisms of glyphosate resistance in *Lolium multiflorum*. Planta 226: 395–404.

Powles SB. 2010. Gene amplification delivers glyphosate-resistant weed evolution. Proceedings of the National Academy of Sciences 107: 955–956.

Pretty J. 2018. Intensification for redesigned and sustainable agricultural systems. Science 362: eaav0294.

Pritchard VL, Mäkinen H, Vähä J-P, Erkinaro J, Orell P, Primmer CR. 2018. Genomic signatures of fine-scale local selection in Atlantic salmon suggest involvement of sexual maturation, energy homeostasis and immune defence-related genes. Molecular Ecology 27: 2560–2575.

Purcell S, Neale B, Todd-Brown K, Thomas L, Ferreira MAR, Bender D, Maller J, Sklar P, Bakker PIW, Daly MJ et al. 2007. PLINK: a tool set for whole-genome association and population-based linkage analyses. The American Journal of Human Genetics 81: 559–575.

Rochette NC, Rivera-Colón AG, Catchen JM. 2019. Stacks 2: analytical methods for paired-end sequencing improve RADseq-based population genomics. Molecular Ecology 28: 4737–4754.

Salas RA, Dayan FE, Pan Z, Watson SB, Dickson JW, Scott RC, Burgos NR. 2012. EPSPS gene amplification in glyphosate-resistant Italian ryegrass (*Lolium perenne ssp. multiflorum*) from Arkansas. Pest Management Science 68: 1223–1230.

Sammons RD, Gaines TA. 2014. Glyphosate resistance: state of knowledge. Pest Management Science 70: 1367–1377.

Schmittgen TD, Livak KJ. 2008. Analyzing real-time PCR data by the comparative CT method. Nature Protocols 3: 1101–1108.

Schroeder J, Barrett M, Shaw DR, Asmus AB, Coble H, Ervin D, Jussaume RA, Owen MDK, Burke I, Creech CF et al. 2018. Managing wicked herbicide-resistance: lessons from the field. Weed Technology 32: 475–488.

Shaner DL, Nadler-Hassar T, Henry WB, Koger CH. 2005. A rapid in vivo shikimate accumulation assay with excised leaf discs. Weed Science 53: 769–774.

Shipley PR, Messinger JJ, Decker AM. 1992. Conserving residual corn fertilizer nitrogen with winter cover crops. Agronomy Journal 84: 869–876.

Stanke M, Keller O, Gunduz I, Hayes A, Waack S, Morgenstern B. 2006. AUGUSTUS: ab initio prediction of alternative transcripts. Nucleic Acids Research 34:, W435–W439.

Steinrücken HC, Amrhein N. 1984. 5-Enolpyruvylshikimate-3-phosphate synthase of Klebsiella pneumoniae. European Journal of Biochemistry 143: 351–357.

Suzukawa AK, Bobadilla LK, Mallory-Smith C, Brunharo CACG. 2021. Non-target-site resistance in *Lolium spp.* globally: a review. Frontiers in Plant Science 11: e609209.

United Nations. 2019. World population Prospects 2019. https://population.un.org/wpp/Graphs/DemographicProfiles/Line/900.

Wickham H. 2016. ggplot2: elegant graphics for data analysis.. New York: Springer-Verlag.

