## Supporting information for "Evidence of multiple origins of glyphosate resistance evolution in *Lolium multiflorum*"

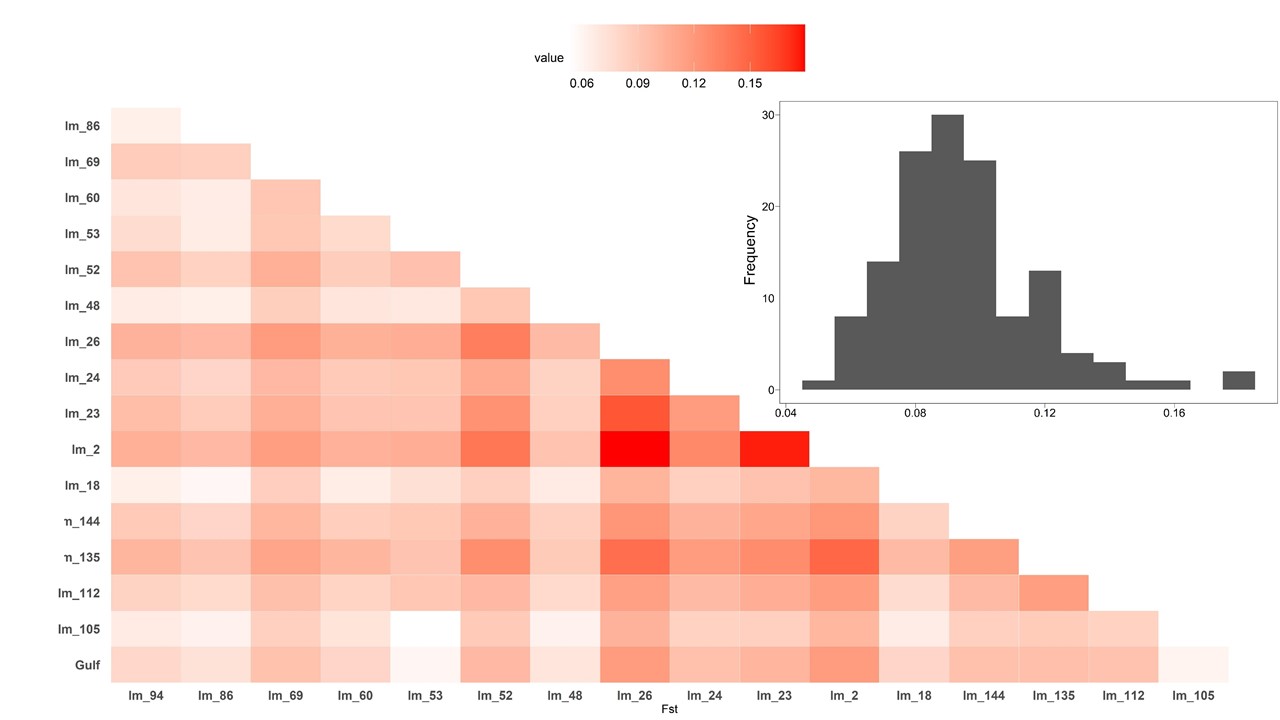

Figure S1. Heatmap of pairwise F_ST_ among glyphosate-resistant and –susceptible *L. multiflorum* populations. Annual ryegrass variety Gulf was included in the analysis. Histogram with distribution of pairwise F_ST_ values from heatmap (mean = 0.09, median=0.09).

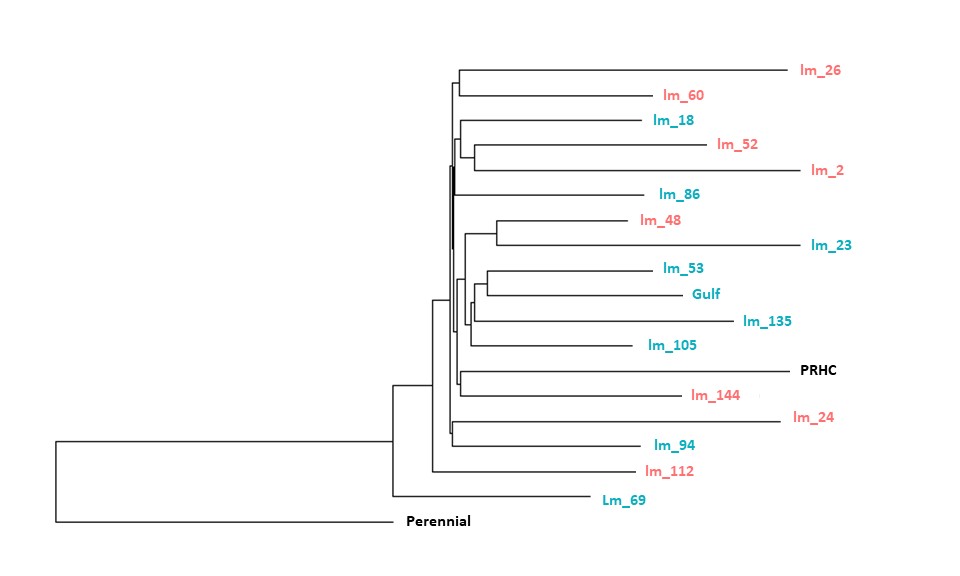

Figure S2. Neighbor joining tree based on pairwise genetic distances (F_ST_) between all pairs of *L. multiflorum* populations from Oregon, using the perennial *L. perenne* as an outgroup. Red and blue are glyphosate-resistant (R) and –susceptible (S), respectively.

Table S1. Geographical location, resistance phenotype, sample size, and mean accumulated shikimate (± SE) for the populations in this study.

| Population | Lat | Long | Crop | Glyphosate resistance^a^ | Multiple resistance^b^ | N | Shikimate (µg g^-1^ FW) |
| --- | --- | --- | --- | --- | --- | --- | --- |
| lm_2 | 45.553 | -123.115 | Orchardgrass | R | ACCase | 14 | 5.1 ± 1.0 |
| lm_18 | 45.397 | -123.111 | Wheat | S | ALS | 16 | 60.8 ± 2.2 |
| lm_23 | 45.241 | -122.952 | Tall fescue | S | - | 14 | 56.0 ± 3.0 |
| lm_24 | 45.224 | -122.861 | Orchardgrass | R | ACCase | 16 | 18.8 ± 5.9 |
| lm_26 | 45.218 | -122.993 | Orchardgrass | R | ACCase | 14 | 4.0 ± 0.9 |
| lm_48 | 45.006 | -122.852 | Wheat | R | - | 16 | 12.0 ± 5.3 |
| lm_52 | 44.955 | -122.884 | Orchardgrass | R | ALS, VLCFA | 16 | 4.1 ± 1.0 |
| lm_53 | 44.922 | -122.735 | Tall fescue | S | - | 16 | 50.1 ± 5.5 |
| lm_60 | 44.884 | -123.257 | Tree crop | R | PSI | 16 | 9.3 ± 3.4 |
| lm_69 | 44.812 | -122.715 | Tall fescue | S | - | 16 | 59.2 ± 4.5 |
| lm_86 | 44.750 | -123.265 | Tall fescue | S | - | 14 | 56.7 ± 1.9 |
| lm_94 | 44.715 | -122.840 | Tree crop | S | ALS | 15 | 56.4 ± 2.0 |
| lm_105 | 44.521 | -123.337 | Tall fescue | S | - | 16 | 62.5 ± 3.4 |
| lm_112 | 44.462 | -123.236 | Wheat | R | ACCase | 16 | 4.9 ± 1.2 |
| lm_135 | 44.359 | -123.053 | White clover | S | - | 14 | 58.6 ± 2.4 |
| lm_144 | 44.279 | -123.137 | Orchardgrass | R | - | 16 | 4.5 ± 1.1 |
| PRHC | 39.752 | -122.016 |  | R | ALS, ACCase, PSI | 13 | 6.0 ± 1.3 |
| Gulf |  |  |  | S |  | 15 | 60.4 ± 2.5 |
| perennial |  |  |  | S |  | 15 | 56.1 ± 1.9 |

^a^R indicates glyphosate resistant, S indicates glyphosate-susceptible. ^b^Indicates to which other mechanism of herbicide action the population is resistant. ACCase: acetyl CoA carboxylase inhibitors (clethodim, pinoxaden, and quizalofop). ALS: acetolactate synthase inhibitors (mesosulfuron and pyroxsulam). VLCFA: very long-chain fatty acid inhibitors (flufenacet). PSI: photosystem I inhibitors (paraquat). N is the number of individuals analyzed per population. Shikimate is the mean concentration of shikimate obtained. Gulf and perennial are cultivated varieties.

Table S2. Primers designed for copy number variation in *L. multiflorum* populations

| Primer name | Target gene | Sequence | Experiment | Fragment size |
| --- | --- | --- | --- | --- |
| AWF^1^ | EPSPS | 5’- AACAGTGAGGAYGTYCACTACATGCT-3’ | EPSPS Sanger sequencing | 338 |
| AWR | EPSPS | 5’- CGAACAGGTGGGCAMTCAGT CT-3’ |  |  |
| EPSPS_Exp_32_F | EPSPS | 5’-AAA GGA TGC CAA GGA GGA AGT-3’ | *EPSPS* copy number | 68 |
| EPSPS_Exp_99_R | EPSPS | 5’-CCG TCA ATG GCC GCA TCG CA-3’ |  |  |
| 514_F | ALS | 5’-CAGCCGCATGTCTCCATTTG-3’ | Housekeeping for EPSPS and ABCC8 | 123 |
| 648_R | ALS | 5’-CTCCCTTTTCTGCTGCTCCA-3’ |  |  |
| ECHCO_1383F | ABCC8 | 5’-GCACTGGTTCTTCGACTCCA-3’ | ABCC8 Sanger sequencing | 420 |
| ECHCO_1800R | ABCC8 | 5’-CTCCCATGACTGCAGCTTGA-3’ |  |  |
| ECHCO_220F | ABCC8 | 5’-GTAGGTACGCTCTTCTGGGC-3’ | ABCC8 gene expression | 119 |
| ECHCO_338R | ABCC8 | 5’-AACTTGGCCTGGTACCCTTG-3’ |  |  |

^1^Primers for *EPSPS* copy number from Abu-Yeboah et al. (2014)

Table S3. Loci within the top 1% of the F_ST_ distribution in all pairwise comparisons.

| Population | Outlier loci |
| --- | --- |
| Lm_2 | 3802 |
|  | 7354 |
|  | 8585 |
|  | 9285 |
|  | 9720 |
|  | 9746 |
|  | 10059 |
|  | 13671 |
|  | 17033 |
|  | 20079 |
|  | 21641 |
|  | 23325 |
|  | 23714 |
|  | 24439 |
|  | 27507 |
| Lm_24 | 19244 |
|  | 15877 |
| Lm_26 | 9144 |
|  | 23899 |
| Lm_60 | 18559 |
|  | 12669 |
|  | 16721 |
|  | 23476 |
| Lm_112 | 3122 |
| Lm_144 | 23923 |

Table S4. Annotations of contigs with SNP’s exhibiting F_ST_ of 1. Contigs were subjected to AUGUSTUS analysis to predict coding proteins, and coding proteins were annotated with Blast2GO 5.

| Description | Length | e-Value | GO IDs | GO Names |
| --- | --- | --- | --- | --- |
| hypothetical protein D1007_44704 | 110 | 3.47E-12 |  |  |
| hypothetical protein D1007_32922 | 124 | 1.23E-15 |  |  |
| putative avenin-like a precursor | 125 | 7.34E-27 | F:GO:0045735 | F:nutrient reservoir activity |
| zinc finger protein 2-like | 138 | 3.22E-13 |  |  |
| predicted protein | 140 | 1.40E-45 | P:GO:0006412; F:GO:0003735; C:GO:0005840 | P:translation; F:structural constituent of ribosome; C:ribosome |
| hypothetical protein BRADI_1g20045v3 | 148 | 3.88E-22 |  |  |
| unnamed protein product | 154 | 9.73E-61 |  |  |
| predicted protein | 157 | 6.38E-75 |  |  |
| Hydroxyethylthiazole kinase | 165 | 4.32E-17 | F:GO:0003676; F:GO:0004523 | F:nucleic acid binding; F:RNA-DNA hybrid ribonuclease activity |
| zinc finger protein 4-like | 167 | 1.19E-22 |  |  |
| uncharacterized protein LOC109774714 | 227 | 1.43E-32 |  |  |
| hypothetical protein SEVIR_2G255500v2 | 241 | 7.32E-11 |  |  |
| predicted protein | 242 | 8.96E-49 |  |  |
| lamin-like protein | 257 | 1.42E-34 | F:GO:0009055 | F:electron transfer activity |
| non-specific lipid-transfer protein C6-like | 257 | 4.69E-36 |  |  |
| 40S ribosomal protein S3a | 264 | 3.12E-163 | P:GO:0006412; F:GO:0003735; C:GO:0005840 | P:translation; F:structural constituent of ribosome; C:ribosome |
| hypothetical protein D1007_28468 | 277 | 1.74E-27 |  |  |
| retrotransposon protein, putative, unclassified | 283 | 7.23E-77 | F:GO:0003676; F:GO:0004523 | F:nucleic acid binding; F:RNA-DNA hybrid ribonuclease activity |
| unnamed protein product | 285 | 2.29E-89 | F:GO:0003676 | F:nucleic acid binding |
| Auxin-induced protein 5NG4 | 286 | 1.94E-70 |  |  |
| putative nuclease HARBI1 | 287 | 2.10E-111 | F:GO:0016788 | F:hydrolase activity, acting on ester bonds |
| acidic endochitinase SE2-like | 299 | 0 | P:GO:0005975 | P:carbohydrate metabolic process |
| uncharacterized protein LOC119271459 | 299 | 1.15E-62 |  |  |
| mannose/glucose-specific lectin-like | 304 | 6.04E-156 | F:GO:0030246 | F:carbohydrate binding |
| Hypothetical predicted protein | 308 | 1.40E-14 |  |  |
| predicted protein | 308 | 1.92E-94 |  |  |
| predicted protein | 317 | 1.48E-122 |  |  |
| retrotransposon protein, putative, unclassified | 318 | 5.38E-75 |  |  |
| GDSL esterase/lipase APG-like | 321 | 4.45E-137 | F:GO:0016788 | F:hydrolase activity, acting on ester bonds |
| predicted protein | 321 | 1.49E-65 |  |  |
| serine/threonine-protein kinase PBL27-like | 333 | 5.71E-129 | P:GO:0006468; F:GO:0004672; F:GO:0005524 | P:protein phosphorylation; F:protein kinase activity; F:ATP binding |
| hypothetical protein CFC21_073247 | 335 | 7.62E-11 |  |  |
| Protein kinase domain superfamily protein | 341 | 1.53E-65 |  |  |
| retrotransposon unclassified | 341 | 4.97E-85 | F:GO:0003676; F:GO:0004523 | F:nucleic acid binding; F:RNA-DNA hybrid ribonuclease activity |
| probable chromo domain-containing protein LHP1 | 355 | 1.15E-140 | P:GO:0006342 | P:chromatin silencing |
| ML domain protein isoform X2 | 361 | 2.99E-60 |  |  |
| lamin-like protein | 371 | 5.58E-49 | F:GO:0009055 | F:electron transfer activity |
| serine/threonine-protein kinase PBL27-like | 376 | 1.48E-123 | P:GO:0006468; F:GO:0004672; F:GO:0005524 | P:protein phosphorylation; F:protein kinase activity; F:ATP binding |
| probable protein phosphatase 2C 77 | 384 | 1.98E-130 | P:GO:0006470; F:GO:0004722 | P:protein dephosphorylation; F:protein serine/threonine phosphatase activity |
| ATP-citrate synthase alpha chain protein 3 | 391 | 0 | F:GO:0005524 | F:ATP binding |
| putative reverse transcriptase | 402 | 1.48E-87 |  |  |
| predicted protein | 410 | 0 |  |  |
| O-methyltransferase ZRP4 | 420 | 0 | F:GO:0008171; F:GO:0046983 | F:O-methyltransferase activity; F:protein dimerization activity |
| predicted protein | 435 | 8.37E-100 | F:GO:0005515 | F:protein binding |
| unnamed protein product | 435 | 3.70E-47 |  |  |
| glutamate receptor 2.8-like | 439 | 0 | F:GO:0015276; C:GO:0016020 | F:ligand-gated ion channel activity; C:membrane |
| predicted protein | 444 | 0 |  |  |
| probable glycosyltransferase 3 | 445 | 0 | F:GO:0016757; C:GO:0016021 | F:transferase activity, transferring glycosyl groups; C:integral component of membrane |
| predicted protein | 447 | 1.17E-124 |  |  |
| BTB/POZ domain-containing protein At3g05675 | 455 | 0 | F:GO:0005515 | F:protein binding |
| aspartic proteinase nepenthesin-1-like | 456 | 7.31E-139 |  |  |
| Hypothetical predicted protein | 462 | 4.23E-31 |  |  |
| unnamed protein product | 465 | 7.92E-36 |  |  |
| uncharacterized protein LOC109767293, partial | 469 | 1.21E-46 |  |  |
| wall-associated receptor kinase-like 6 | 470 | 0 | P:GO:0006468; F:GO:0004672; F:GO:0005509; F:GO:0005524 | P:protein phosphorylation; F:protein kinase activity; F:calcium ion binding; F:ATP binding |
| Anthranilate synthase component I-1, chloroplastic | 477 | 2.14E-34 |  |  |
| uncharacterized protein LOC119315563 | 479 | 4.30E-94 | F:GO:0005515 | F:protein binding |
| unnamed protein product | 484 | 1.11E-84 |  |  |
| chromatin assembly factor 1 subunit p90-like | 490 | 1.66E-27 |  |  |
| wall-associated receptor kinase 2-like isoform X1 | 495 | 0 | F:GO:0030247 | F:polysaccharide binding |
| Nitrate transporter | 501 | 2.37E-75 |  |  |
| Transposon protein, putative, Mutator sub-class | 501 | 8.16E-24 |  |  |
| glutamate receptor 2.8-like | 507 | 0 |  |  |
| lamin-like protein | 513 | 4.39E-45 | F:GO:0009055 | F:electron transfer activity |
| uncharacterized protein LOC119332870 | 514 | 3.89E-48 |  |  |
| uncharacterized protein LOC109739135 | 516 | 0 |  |  |
| predicted protein | 520 | 0 | F:GO:0005506; F:GO:0016705; F:GO:0020037 | F:iron ion binding; F:oxidoreductase activity, acting on paired donors, with incorporation or reduction of molecular oxygen; F:heme binding |
| dna replication licensing factor mcm2 | 526 | 2.47E-56 |  |  |
| unnamed protein product | 526 | 5.76E-71 | F:GO:0003676; F:GO:0004523 | F:nucleic acid binding; F:RNA-DNA hybrid ribonuclease activity |
| putative methionyl-tRNA synthetase | 526 | 3.80E-23 |  |  |
| Anthranilate synthase component I-1, chloroplastic | 529 | 3.34E-40 | F:GO:0003676; F:GO:0008270 | F:nucleic acid binding; F:zinc ion binding |
| hypothetical protein PAHAL_3G517500 | 529 | 1.67E-54 |  |  |
| predicted protein | 530 | 0 | F:GO:0005506; F:GO:0016705; F:GO:0020037 | F:iron ion binding; F:oxidoreductase activity, acting on paired donors, with incorporation or reduction of molecular oxygen; F:heme binding |
| membrane magnesium transporter | 542 | 2.46E-40 |  |  |
| predicted protein | 542 | 0 | F:GO:0005506; F:GO:0016705; F:GO:0020037 | F:iron ion binding; F:oxidoreductase activity, acting on paired donors, with incorporation or reduction of molecular oxygen; F:heme binding |
| hypothetical protein CFC21_070598 | 545 | 0 |  |  |
| predicted protein | 545 | 6.69E-100 | P:GO:0006355; F:GO:0003677 | P:regulation of transcription, DNA-templated; F:DNA binding |
| putative AP2 protein | 549 | 1.68E-16 | P:GO:0006355; F:GO:0003677; F:GO:0003700 | P:regulation of transcription, DNA-templated; F:DNA binding; F:DNA-binding transcription factor activity |
| uncharacterized protein LOC119363449 | 551 | 7.85E-29 |  |  |
| hypothetical protein BRADI_1g46192v3 | 552 | 4.04E-45 |  |  |
| hypothetical protein D1007_17136 | 563 | 2.94E-28 |  |  |
| pre-mRNA-splicing factor CWC25 homolog | 569 | 0 |  |  |
| pentatricopeptide repeat-containing protein At2g45350, chloroplastic-like | 569 | 0 | F:GO:0005515 | F:protein binding |
| Glutamate receptor 2.7 | 572 | 1.05E-177 | F:GO:0004970; C:GO:0016020 | F:ionotropic glutamate receptor activity; C:membrane |
| zinc finger MYM-type protein 1-like | 581 | 3.78E-37 | F:GO:0046983 | F:protein dimerization activity |
| retrotransposon protein, putative, Ty3-gypsy subclass | 583 | 0 | F:GO:0003676; F:GO:0004523 | F:nucleic acid binding; F:RNA-DNA hybrid ribonuclease activity |
| retrotransposon protein, putative, unclassified | 587 | 7.37E-18 |  |  |
| cytochrome P450 72A15-like | 592 | 0 | F:GO:0005506; F:GO:0016705; F:GO:0020037 | F:iron ion binding; F:oxidoreductase activity, acting on paired donors, with incorporation or reduction of molecular oxygen; F:heme binding |
| protein FRIGIDA | 594 | 4.33E-43 |  |  |
| CCR4-NOT transcription complex subunit 11-like isoform X1 | 596 | 0 | C:GO:0030014 | C:CCR4-NOT complex |
| putative LRR receptor-like serine/threonine-protein kinase | 606 | 0 | F:GO:0005515 | F:protein binding |
| probable inactive leucine-rich repeat receptor kinase XIAO | 616 | 0 | F:GO:0005515 | F:protein binding |
| putative retrotransposon protein | 619 | 1.26E-61 |  |  |
| hypothetical protein D1007_43406 | 627 | 2.05E-139 | P:GO:0006313; F:GO:0003677; F:GO:0004803; F:GO:0008270 | P:transposition, DNA-mediated; F:DNA binding; F:transposase activity; F:zinc ion binding |
| transmembrane 9 superfamily member 9-like | 636 | 0 | C:GO:0016021 | C:integral component of membrane |
| E3 ubiquitin-protein ligase RGLG2-like | 642 | 0 |  |  |
| uncharacterized protein LOC119297384 | 650 | 1.57E-105 |  |  |
| glutamate receptor 2.8-like | 654 | 4.69E-130 | F:GO:0015276; C:GO:0016020 | F:ligand-gated ion channel activity; C:membrane |
| predicted protein | 656 | 0 |  |  |
| Alpha-galactosidase | 665 | 8.58E-46 |  |  |
| uncharacterized protein LOC112269893 | 666 | 5.15E-36 |  |  |
| retrotransposon gag domain-containing protein | 678 | 9.73E-59 |  |  |
| uncharacterized protein LOC119308629 | 684 | 0 |  |  |
| retrotransposon protein, putative, unclassified | 691 | 1.54E-51 |  |  |
| retrotransposon protein, putative, unclassified | 696 | 8.67E-75 |  |  |
| putative methionyl-tRNA synthetase | 705 | 1.62E-32 |  |  |
| NADPH--cytochrome P450 reductase-like | 714 | 0 | F:GO:0010181; F:GO:0016491 | F:FMN binding; F:oxidoreductase activity |
| transposon protein, putative, CACTA, En/Spm sub-class | 730 | 2.79E-77 |  |  |
| uncharacterized protein LOC109742836 | 738 | 0 | C:GO:0031011 | C:Ino80 complex |
| uncharacterized protein LOC119358248 | 740 | 0 | F:GO:0008270 | F:zinc ion binding |
| predicted protein | 752 | 6.72E-162 |  |  |
| B3 domain-containing protein | 753 | 7.59E-45 |  |  |
| Anthranilate synthase component I-1, chloroplastic | 753 | 1.06E-44 |  |  |
| hypothetical protein D1007_05146 | 756 | 6.00E-14 |  |  |
| PREDICTED: uncharacterized protein LOC102712212 isoform X2 | 759 | 1.52E-40 |  |  |
| proline transporter 1-like | 771 | 4.31E-64 |  |  |
| Retrovirus-related Pol polyprotein from transposon 297 family | 785 | 5.89E-46 |  |  |
| hypothetical protein CFC21_001384 | 795 | 0 |  |  |
| putative pentatricopeptide repeat-containing protein At1g13630 | 795 | 0 | F:GO:0005515 | F:protein binding |
| putative gag-pol precursor -orf2 | 811 | 3.85E-123 | F:GO:0003676 | F:nucleic acid binding |
| transposon protein, putative, CACTA, En/Spm sub-class | 826 | 1.53E-51 |  |  |
| putative pentatricopeptide repeat-containing protein At3g25970 | 859 | 0 | F:GO:0005515 | F:protein binding |
| uncharacterized protein LOC104584952 | 860 | 1.44E-62 |  |  |
| probable protein phosphatase 2C 66 | 864 | 0 | F:GO:0003677; F:GO:0016791 | F:DNA binding; F:phosphatase activity |
| cyclic nucleotide-gated ion channel 17-like | 865 | 0 | P:GO:0006811; P:GO:0055085; F:GO:0005216; C:GO:0016020 | P:ion transport; P:transmembrane transport; F:ion channel activity; C:membrane |
| putative ribonuclease H protein | 892 | 5.84E-23 |  |  |
| dof zinc finger protein | 898 | 2.92E-74 | P:GO:0006355; F:GO:0003677 | P:regulation of transcription, DNA-templated; F:DNA binding |
| gag polyprotein | 916 | 6.25E-27 |  |  |
| retrotransposon unclassified | 917 | 1.24E-146 |  |  |
| retrotransposon protein, putative, unclassified | 932 | 0 | F:GO:0003676; F:GO:0004523 | F:nucleic acid binding; F:RNA-DNA hybrid ribonuclease activity |
| eukaryotic translation initiation factor 3 subunit C | 933 | 0 | P:GO:0006413; F:GO:0003743; F:GO:0031369; C:GO:0005852 | P:translational initiation; F:translation initiation factor activity; F:translation initiation factor binding; C:eukaryotic translation initiation factor 3 complex |
| glutamate receptor 2.8-like | 940 | 0 | F:GO:0015276; C:GO:0016020 | F:ligand-gated ion channel activity; C:membrane |
| cytochrome P450 72A15-like | 949 | 0 | F:GO:0005506; F:GO:0016705; F:GO:0020037 | F:iron ion binding; F:oxidoreductase activity, acting on paired donors, with incorporation or reduction of molecular oxygen; F:heme binding |
| protein FAR1-RELATED SEQUENCE 5-like isoform X1 | 957 | 2.33E-75 | F:GO:0003676; F:GO:0008270 | F:nucleic acid binding; F:zinc ion binding |
| retrotransposon protein, putative, unclassified | 961 | 4.66E-178 | F:GO:0003676; F:GO:0004523 | F:nucleic acid binding; F:RNA-DNA hybrid ribonuclease activity |
| peroxisome biogenesis protein 5 isoform X1 | 963 | 0 | F:GO:0005515 | F:protein binding |
| Putative retroelement | 993 | 9.00E-133 | P:GO:0015074; F:GO:0003676 | P:DNA integration; F:nucleic acid binding |
| hypothetical protein | 1000 | 3.23E-29 | C:GO:0016020; C:GO:0016021 | C:membrane; C:integral component of membrane |
| G-type lectin S-receptor-like serine/threonine-protein kinase LECRK3 | 1018 | 0 | P:GO:0006468; F:GO:0004672; F:GO:0005524 | P:protein phosphorylation; F:protein kinase activity; F:ATP binding |
| Endoglucanase 3 | 1046 | 3.08E-71 |  |  |
| putative cytokinin riboside 5'-monophosphate phosphoribohydrolase LOGL10 | 1052 | 1.17E-46 |  |  |
| predicted protein | 1053 | 2.51E-107 | F:GO:0005515 | F:protein binding |
| glutamate receptor 2.8-like | 1081 | 0 | F:GO:0015276; C:GO:0016020 | F:ligand-gated ion channel activity; C:membrane |
| unnamed protein product | 1092 | 8.33E-58 | F:GO:0016788 | F:hydrolase activity, acting on ester bonds |
| thiamine phosphate phosphatase-like protein | 1135 | 2.98E-132 | F:GO:0016791 | F:phosphatase activity |
| F-box protein At5g07610-like | 1175 | 3.62E-62 |  |  |
| probable protein phosphatase 2C 70 isoform X1 | 1188 | 2.64E-161 | P:GO:0006470; F:GO:0004722 | P:protein dephosphorylation; F:protein serine/threonine phosphatase activity |
| uncharacterized protein LOC100830937 isoform X1 | 1224 | 9.19E-52 |  |  |
| ARF guanine-nucleotide exchange factor GNOM-like | 1234 | 0 | P:GO:0032012; F:GO:0005085 | P:regulation of ARF protein signal transduction; F:guanyl-nucleotide exchange factor activity |
| protein SUPPRESSOR OF GENE SILENCING 3 homolog | 1314 | 0 | P:GO:0031047; P:GO:0051607 | P:gene silencing by RNA; P:defense response to virus |
| putative polyprotein | 1365 | 0 | P:GO:0015074; F:GO:0003676 | P:DNA integration; F:nucleic acid binding |
| protein NRT1/ PTR FAMILY 5.10-like | 1371 | 0 | P:GO:0055085; F:GO:0022857; C:GO:0016020 | P:transmembrane transport; F:transmembrane transporter activity; C:membrane |
| Protein kinase domain superfamily protein | 1410 | 0 | P:GO:0055085; C:GO:0016021 | P:transmembrane transport; C:integral component of membrane |
| wall-associated receptor kinase 2-like | 1412 | 0 | P:GO:0006468; F:GO:0004672; F:GO:0005524; F:GO:0030247 | P:protein phosphorylation; F:protein kinase activity; F:ATP binding; F:polysaccharide binding |
| retrotransposon protein, putative, Ty1-copia subclass | 1438 | 4.98E-41 | P:GO:0006810; P:GO:0009987; F:GO:0005488 | P:transport; P:cellular process; F:binding |
| predicted protein | 1520 | 0 |  |  |
| predicted protein | 1552 | 9.58E-111 |  |  |
| hypothetical protein TRIUR3_29628 | 1555 | 0 |  |  |
| retrotransposon protein, putative, unclassified | 1626 | 0 | P:GO:0015074; F:GO:0003676 | P:DNA integration; F:nucleic acid binding |
| hypothetical protein CFC21_064604 | 1652 | 0 |  |  |
| tRNA-specific adenosine deaminase TAD3-like isoform X1 | 1697 | 7.07E-94 | F:GO:0003824 | F:catalytic activity |
| retrotransposon protein, putative, Ty3-gypsy subclass | 1755 | 6.15E-28 | P:GO:0015074; P:GO:0090502; F:GO:0003676; F:GO:0004523 | P:DNA integration; P:RNA phosphodiester bond hydrolysis, endonucleolytic; F:nucleic acid binding; F:RNA-DNA hybrid ribonuclease activity |
| putative disease resistance RPP13-like protein 3 | 1795 | 0 | F:GO:0008270; F:GO:0043531 | F:zinc ion binding; F:ADP binding |
| Putative polyprotein | 1803 | 0 | P:GO:0015074; F:GO:0003676 | P:DNA integration; F:nucleic acid binding |
| retrotransposon protein, putative, unclassified | 1846 | 0 | F:GO:0003676; F:GO:0008270 | F:nucleic acid binding; F:zinc ion binding |
| uncharacterized protein LOC104583208 | 1861 | 3.28E-113 | P:GO:0015074; F:GO:0003676; F:GO:0004523 | P:DNA integration; F:nucleic acid binding; F:RNA-DNA hybrid ribonuclease activity |
| Protein kinase domain superfamily protein | 2126 | 0 |  |  |
| DNA-directed RNA polymerase V subunit 1-like | 2258 | 0 | P:GO:0006351; F:GO:0003677; F:GO:0003899 | P:transcription, DNA-templated; F:DNA binding; F:DNA-directed 5'-3' RNA polymerase activity |
| serine/threonine-protein kinase EDR1 | 2272 | 0 | P:GO:0006468; F:GO:0004672; F:GO:0005524; F:GO:0016788 | P:protein phosphorylation; F:protein kinase activity; F:ATP binding; F:hydrolase activity, acting on ester bonds |
| retrotransposon protein, putative, Ty1-copia subclass | 2383 | 4.51E-158 | P:GO:0015074; F:GO:0003676 | P:DNA integration; F:nucleic acid binding |
